# Duffy Antigen Expression in Erythroid Bone Marrow Precursor Cells of Genotypically Duffy Negative Individuals

**DOI:** 10.1101/508481

**Authors:** Célia Dechavanne, Sebastien Dechavanne, Sylvain Metral, Brooke Roeper, Sushma Krishnan, Rich Fong, Seth Bennett, Lenore Carias, Edwin Chen, Nichole D. Salinas, Anil Ghosh, Niraj H. Tolia, Philip G. Woost, James W. Jacobberger, Yves Colin, Benoit Gamain, Christopher L. King, Peter A. Zimmerman

## Abstract

The gene encoding the Duffy blood group protein (Fy, CD234; additional designations Duffy Antigen Receptor of Chemokines [DARC] and Atypical Chemokine Receptor 1 [ACKR1]) is characterized by a SNP in a GATA-1 transcription factor binding site associated with the erythrocyte silent (ES) phenotype. *FY*^*ES*^ homozygous people are viewed to be highly resistant to blood stage infection with *Plasmodium vivax*. Increasingly, however, studies are reporting *P. vivax* infections in Fy-negative individuals across malarious African countries where *FY*^*ES*^ approaches genetic fixation. This suggests that *P. vivax* has evolved a Fy-independent RBC invasion pathway, or that the GATA-1 SNP does not abolish Fy expression. Here, we tested the second hypothesis through binding studies to erythroid lineage cells using recombinant *P. vivax* Duffy binding protein, the parasite’s invasion ligand and Fy6-specific antibodies. We first observed variable Fy expression on circulating RBCs, irrespective of *FY* genotype; *FY*^*ES*^ RBCs were periodically Fy-positive. Furthermore, during the *in vitro* erythroid differentiation of CD34+ cells and on *ex vivo* bone marrow samples, we observed Fy expression on erythroid precursor cells from *FY*^*ES*^ people. Finally, the Fy6-specific nanobody, CA111 was used to capture Fy from the surface of *FY*^*ES*^ RBCs. Our findings reveal that the GATA-1 SNP does not fully abolish Fy expression and provide insight on potential susceptibility of Fy-negative people to vivax malaria.

**Significance:** Duffy blood group negativity results from a single nucleotide polymorphism (SNP) in the gene promoter, and reaches genetic fixation in many African ethnicities. Because the Duffy protein (Fy) is an important contact point during *Plasmodium vivax* human red blood cell invasion, Fy-negativity is considered to confer resistance to *P. vivax* malaria. With recent studies in African countries reporting *P. vivax* infection in Fy-negative people, we studied Fy expression across erythroid development. Here we report that the *FY* promoter SNP does not abolish Fy protein expression in erythroid progenitors developing in the bone marrow. These results further emphasizes the importance of reticulocytes as targets for *P. vivax* blood stage infection and propose a mechanism for *P. vivax* infections in Fy-negative people.

## Introduction

Duffy gene (*FY*) polymorphism is prominent historically in many corners of biomedical investigation. In 1968, Donahue et al used Fya/ Fyb serological inheritance patterns and human chromosome 1 centromeric staining features to make *FY* the first gene mapped to a human autosome (1). Similarly, the Duffy blood group protein (Fy) (2) was the first identified member of the diverse seven transmembrane chemokine receptor family, albeit with a number of interesting structural and functional differences that make its biology unique (2). Fy expression on both erythroid and non-erythroid cells (2) modulates homeostatic levels and gradients of chemokines between blood circulation (3-7) and tissues (8-12) underlying immune responses and an emerging array of pathologies.

The decades-long observations that Africans and African-Americans, who are for the most part Fy-negative, were highly resistant to *Plasmodium vivax* blood stage infection (13-18) suggested that interaction with the Fy protein was required for parasite invasion of human red blood cells (RBCs) (19). With gene sequencing, a single nucleotide polymorphism (SNP) in the *FY* gene GATA-1 transcription factor binding site has been shown to block *FY* erythroid expression (20, 21) and homozygosity underlies the “Fy erythrocyte silent” (herein, Fy-negative) phenotype characterized by standard serological methods (20). *In vitro* studies using Fy-dependent, *P. knowlesi* (22, 23), have gone on to show that while the merozoite is able to orient apically to the RBC surface, the mechanism fails to establish parasite-Fy-negative RBC commitment. Therefore, onward formation of a tight mobile junction needed to infect the RBC was inhibited, and invasion was blocked (24). This background has inextricably linked Duffy blood group genetics and malaria.

Recent studies now report *P. vivax* infections in Fy-negative individuals from many malarious countries across Africa (25). These observations lead to at least two hypotheses. First, the parasite’s Duffy binding protein (PvDBP) may not interact with Fy, exclusively. Second, constitutive low level or periodic Fy expression may occur in Fy-negative people and facilitate blood stage infection.

Previous studies have demonstrated a gene dosage effect of the *FY* GATA-1 SNP that reduces erythrocyte surface expression of Fy^a^ and Fy^b^ by 50% in heterozygous individuals (21, 26), and others have identified additional coding region SNPs associated with reduced expression of both Fy^a^ (Fy^a^-weak) (27) and Fy^b^ (Fy^b^-weak) (28-30); very rare nonsense SNPs and deletions in the *FY* gene coding region have also been reported (**see text with Supplemental Information (*SI*) Table S1**) (31, 32). Further, Fy is more highly expressed on reticulocytes vs. older RBC (26). With the range of Fy expression phenotypes, here we tested the hypothesis that Fy protein expression may not be fully blocked by the GATA-1 SNP (*FY*^*ES*^ mutation) as demonstrated through previous study designs (33, 34).

## Results

### Binding specificity of CA111

To assess the presence of Fy on RBCs, the camelid single-domain antibody CA111 was used. This VHH or nanobody is specific for the 5-amino acid sequence ^22^FEDVW^26^ within the Fy amino terminal domain, overlapping the linear epitope recognized by the anti-Fy6, 2C3 mouse MoAb (**Supplemental Methods; Text S1**) (35). This is the same region of Fy to which chemokines and PvDBP bind and acts as the receptor for *P. vivax* RBC invasion (35). In specificity assessments, we demonstrate that the anti-Fy6 antibodies CA111 and 2C3, reciprocally inhibit their binding to Fy-positive pbRBCs in competitive binding studies (**Figure S1A**). Additional comparisons among CA111, 2C3, anti-Fya and anti-Fyb show that CA111 demonstrates the strongest relative sensitivity for Fy binding among these antibody reagents (**Figure S1B**). Finally, while *in silico* analysis (BLASTp) showed the Fy ^22^FED<u>V</u>W^26^ and CXCR2 ^11^FED<u>F</u>W^15^ sequence motifs vary at only one position, we demonstrated that CA111 was not able to bind or to pull down CXCR2 (**Figure S2**).

We then tested whether CA111 recognized Fy on *FY*A/*A, FY*B/*B* and *FY*B*^*ES*^*/*B*^*ES*^ RBC. Working on freshly collected, peripheral blood red blood cells (pbRBC), we observed expected binding to pbRBC of *FY*A/*A, FY*B/*B* donors, but also observed unexpected CA111 binding to *FY*B*^*ES*^*/*B*^*ES*^ pbRBC (**Figure 1**). Over three different time points of our initial experiments, the CA111 binding varied from 74.0% to 89.4% in *FY*B/*B* pbRBC, from 77.2% to 90.8% in *FY*A/*A* pbRBC and from 11.5% to 47.6% in *FY*B*^*ES*^*/*B*^*ES*^. This latter result, observed several times and tested in triplicate, was markedly higher than previously observed on *FY*B*^*ES*^*/*B*^*ES*^ donors(35), and suggested that the Fy protein may be expressed periodically on the RBC surface of *FY*B*^*ES*^*/*B*^*ES*^ (Donor #1).

**Figure 1.**
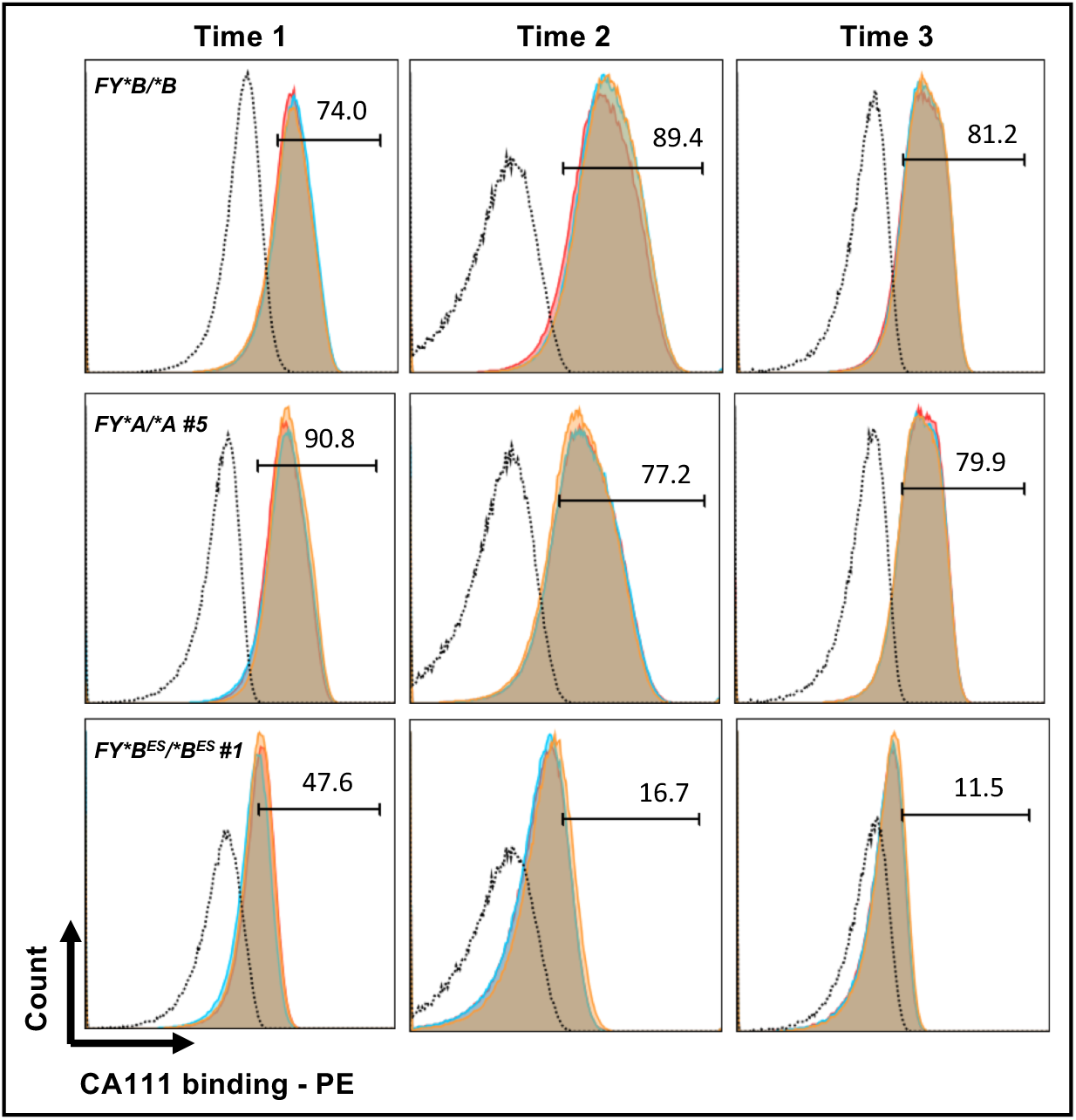
Binding of CA111 on FY*BES/*BES peripheral blood RBC (pbRBC). Direct binding of CA111 (30ug/mL) to pbRBC from *FY*B/*B, FY*A/*A* and *FY*B*^*ES*^*/*B*^*ES*^ (#1) donors. Our preliminary experiments were performed at 3 different time points and in triplicate (histograms in red, orange and blue in overlay). Individual donor samples at each time point served as negative controls represented in each histogram by the dark dashed line. The percentage of cells with CA111 binding (PE: phycoerythrin) above the background established by comparison to negative controls (minus CA111; plus anti-HA mouse antibody; plus anti-mouse-phycoerythrin conjugateted goat antibody) is indicated above the gate determined for positive rPvDBPII binding.

### rPvDBPII binding on *FY*B*^*ES*^*/*B*^*ES*^ peripheral blood RBC

We next tested if rPvDBPII would interact with *FY*B*^*ES*^*/*B*^*ES*^ (Donor #1) pbRBCs. rPvDBPII bound to both *FY*A/*B* and to a lesser extent *FY*B*^*ES*^*/*B*^*ES*^ (Donor #1) pbRBC. As a control, unfolded rPvDBPII failed to bind pbRBC from the same donors (**Figure 2**). rPvDBPII, as well as chemokine binding to Fy have been shown to be sensitive to chymotrypsin treatment of pbRBC(22, 36). Consistent with expectations from these previous studies, we observed that chymotrypsin treatment eliminated rPvDBPII binding to pbRBC from both *FY*A/*B* and *FY*B*^*ES*^*/*B*^*ES*^ donors.

**Figure 2.**
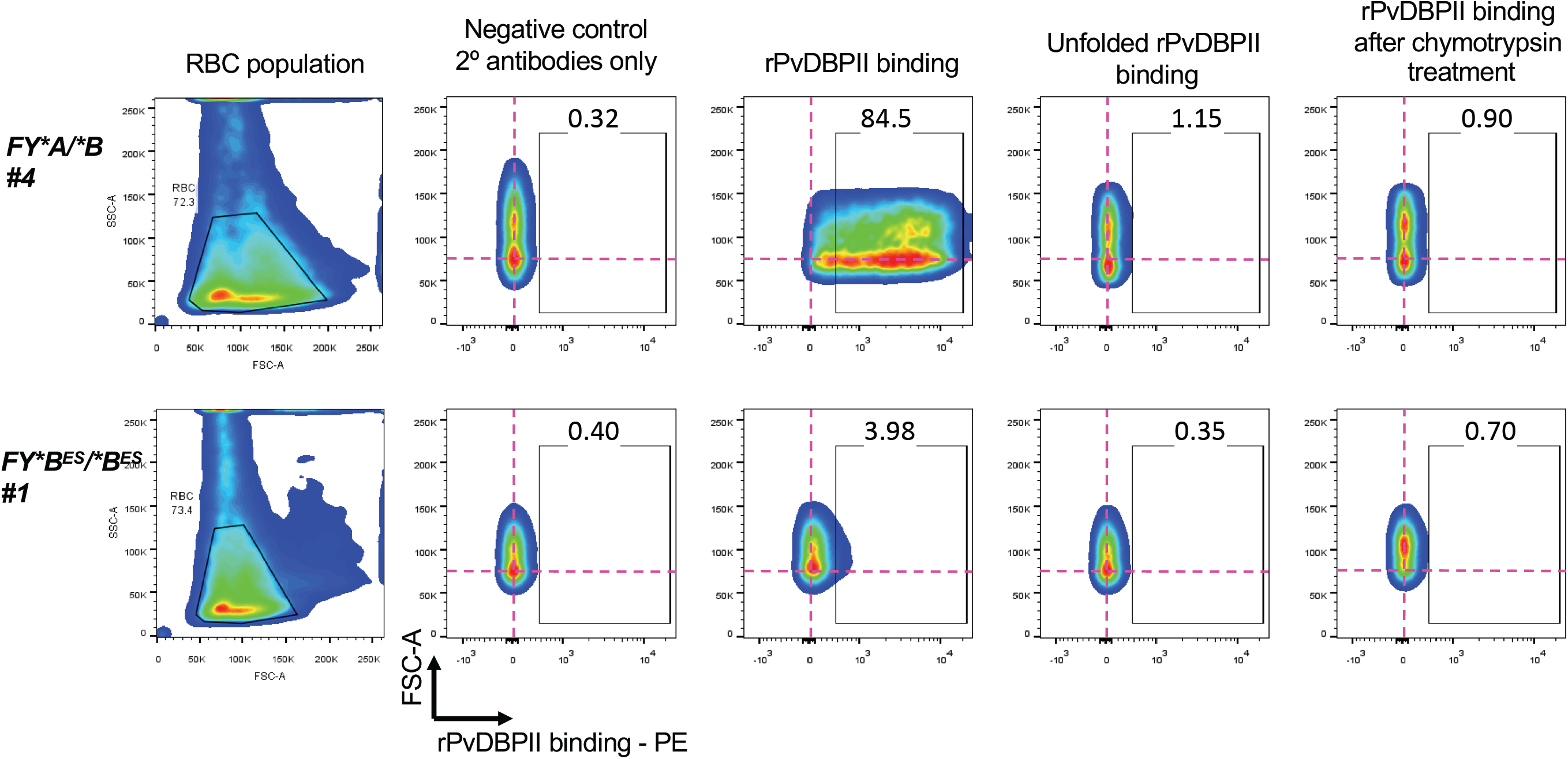
Gating strategy of rPvDBPII binding to pbRBC. The pbRBC population is selected in the forward (FSC-A) and size (SSC-A) scatter plots (left-most panel). The doublets were removed from the analysis (data not shown). The binding of recombinant *Plasmodium vivax* Duffy binding protein, region II (rPvDBPII) on pbRBC is represented by the gated region designated by the open box in each plot. The percentage of pbRBC showing binding to rPvDBPII is indicated above the gate in each experiment. The pink crosshairs show the center of the pbRBC population from the negative control condition (minus rPvDBPII; plus secondary anti-rPvDBPII, rabbit polyclonal antibody; plus anti-rabbit-phycoerythrin-conjugateted, goat antibody) to onward experimental conditions (plus rPvDBPII; plus secondary antibodies). The fixed position of the crosshairs facilitate comparisons between pbRBC population shifts across negative controls and test conditions.

### Specificity of rPvDBPII binding on *FY*B*^*ES*^*/*B*^*ES*^ pbRBC

Given the unexpected interaction of both CA111 and rPvDBPII with *FY*B*^*ES*^*/*B*^*ES*^ pbRBC, we pursued additional experiments to further evaluate the specificity of rPvDBPII binding. We first evaluated the specificity of this interaction by testing rPvDBPII binding to Fy-positive and Fy-negative pbRBC by Western blot analysis (**Figure 3A**).

**Figure 3.**
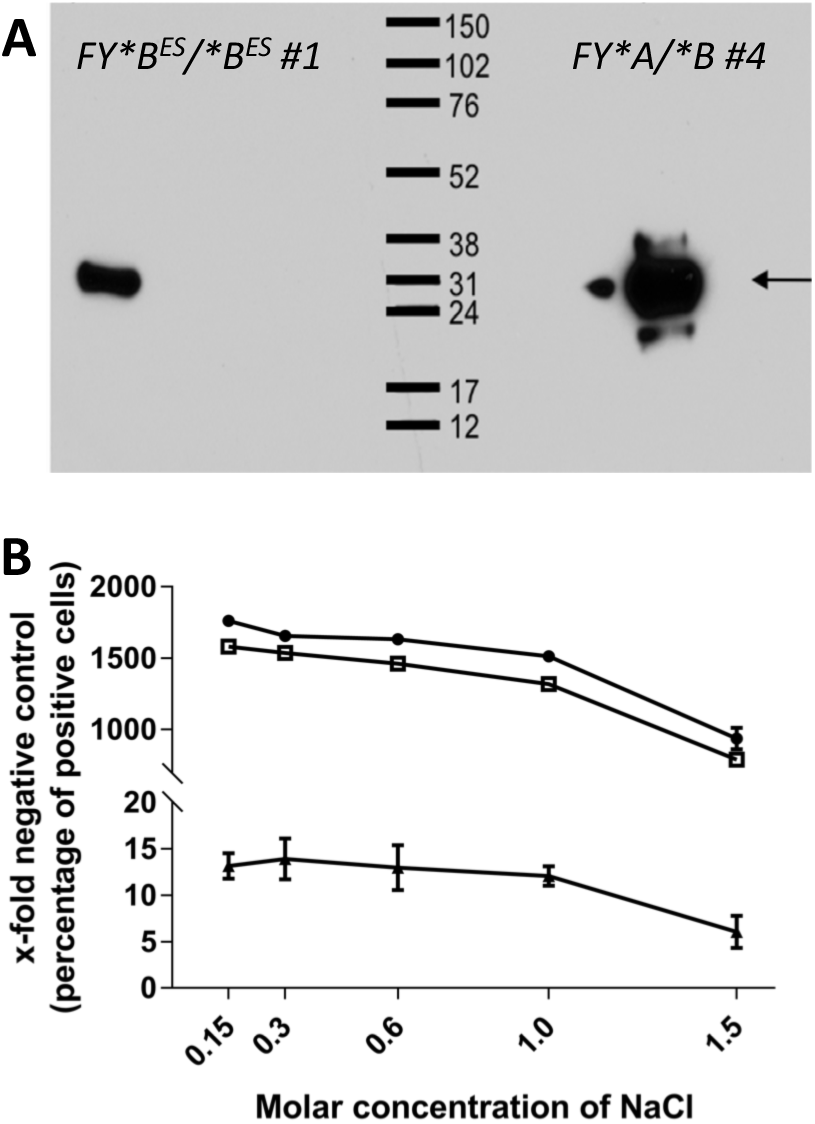
rPvDBPII dissociation and binding to Fy on Fy-positive and Fy-negative pbRBC. Panel A – pbRBC were exposed to rPvDBPII (20 μg/mL) to allow protein interaction and binding with RBC cell-surface proteins (2 hr. at room temperature). Polyclonal anti-PvDBP was used to probe a Western blot of solubilized proteins to determine if the Fy protein was responsible for the rPvDBPII-specific signals observed in flow cytometry experiments and designated by the black arrow. Individual donors for this study included the *FY* genotypes *FY*B*^*ES*^*/*B*^*ES*^ and *FY*A/*B*. The western blot was probed with polyclonal anti-PvDBP and anti-rabbit HRP. For both *FY*B*^*ES*^*/*B*^*ES*^ and *FY*A/*B* genotypes, rPvDBPII bound to a single protein band, with a MW comparable to the Fy protein (35 kD). Panel B - Increasing molar concentrations of NaCl were used to determine if it was possible to dissociate the binding of rPvDBPII from pbRBC. Individual donors for this study included the *FY* genotypes *FY*B/*B, FY*A/*B* and *FY*B*^*ES*^*/*B*^*ES*^. Non-specific binding should be dissociated with a large relative volume (750 μL) of PBS (0.15M of NaCl). No significant reduction in rPvDBPII binding was observed across increasing NaCl concentrations from 0.15M to 1.0M. Only upon addition of 1.5M NaCl was there any significant reduction in rPvDBPII binding. (Pearson Chi^2^ test for median, p=0.014: Pearson Chi^2^=6.0). Each curve represents a different donor: *FY*B/*B* #3 in black circle, *FY*A/*B* #4 in open square, *FY*B*^*ES*^*/*B*^*ES*^ #1 in black triangle.

We exposed both Fy-positive and Fy-negative pbRBC to rPvDBPII, eluted pbRBC bound proteins and then probed a Western blot of these proteins with a polyclonal antibody to PvDBP. Western blot results revealed a single protein band from both *FY*A/*B* and *FY*B*^*ES*^*/*B*^*ES*^ pbRBC (**Figure 3A**); specific detection by the polyclonal anti-PvDBP and the apparent molecular weight of approximately 35kDa were both consistent with detection of the rPvDBPII protein.

To evaluate the relative affinity of rPvDBPII interaction to Fy-positive and Fy-negative pbRBC, we next tested the dissociation of PvDBPII from pbRBC with increasing NaCl concentrations (**Figure 3B**). While we observed that the relative binding of rPvDBPII to *FY*B*^*ES*^*/*B*^*ES*^ pbRBC was markedly lower than that observed for *FY*B/*B* and *FY*A/*B*, dissociation of rPvDBPII from *FY*B/*B* and *FY*A/*B* versus *FY*B*^*ES*^*/*B*^*ES*^ pbRBC was similar across decreasing NaCl concentrations (0.15M, 0.3M, 0.6M and 1M NaCl). At 1.5M NaCl the pbRBC for all *FY* genotypes were lysing and therefore rPvDBPII signal decreased across all samples.

Given consistent demonstration of the chymotrypsin-sensitive nature of the Fy protein(22, 33, 36-38) and the molecular interactions tethered to RBC Fy, we tested the reproducibility of rPvDBPII binding to *FY*B*^*ES*^*/*B*^*ES*^ pbRBC after chymotrypsin treatment. Here, we examined the chymotrypsin sensitivity of rPvDBPII binding for one *FY*A/*B* and *FY*B*^*ES*^*/*B*^*ES*^ (Donor #1) at multiple independent times and results (**Figure 4**) consistently showed that the binding of rPvDBPII to both *FY*A/*B* and *FY*B*^*ES*^*/*B*^*ES*^ pbRBC were significantly higher before than after a chymotrypsin treatment; consistent with results in **Figure 1**, we observed that rPvDBPII binding to Fy varied for the individual donors between different time points.

**Figure 4.**
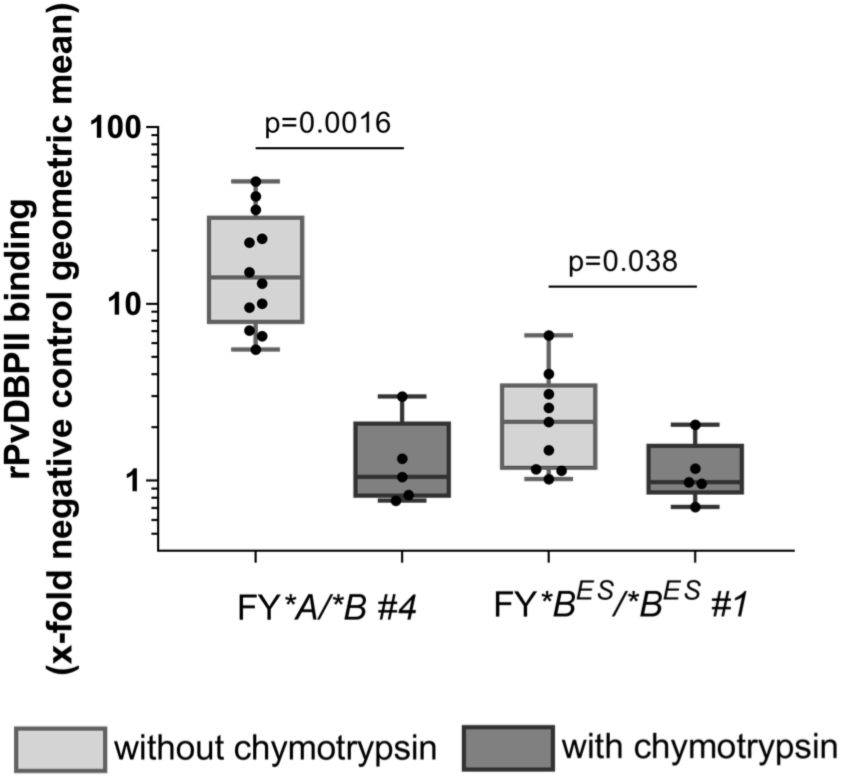
rPvDBPII binding on both Fy-positive and Fy-negative pbRBC is chymotrypsin sensitive.. This experiment assessed the reproducibility of chymotrypsin-sensitive rPvDBPII binding to pbRBC. Individual donors for this study included the *FY* genotypes *FY*A/*B* and *FY*B*^*ES*^*/*B*^*ES*^ using pbRBC without chymotrypsin treatment (light gray boxes) or with chymotrypsin treatment (dark gray boxes); specific individual data points appear as fill black circles. rPvDBPII binding was performed after 5 washes to remove any residual chymotrypsin. Consistent with earlier experiments, individual donor samples served as negative controls (minus rPvDBPII; plus secondary anti-rPvDBPII, rabbit polyclonal antibody; plus anti-rabbit-phycoerythrin conjugateted goat antibody). The measurements for the individual donors were performed at different times. For *FY*A/*B*: without chymotrypsin n=12; with chymotrypsin n=5; for *FY*B*^*ES*^*/*B*^*ES*^: without chymotrypsin n=9; with chymotrypsin n=5. A non-parametric paired Wilcoxon rank-sum test was used to assess the differences between chymotrypsin treatments. *FY*A/*B* z=3,162 (*P-value* = 0.0016); *FY*B*^*ES*^*/*B*^*ES*^ z=2,067 (*P-value* = 0.038).

### rPvDBPII binding on RBC over time

Because we observed variation in CA111 and PvDBPII binding to pbRBC at different time points for both Fy-positive and Fy-negative donors, we examined this variability in greater detail using comparable staining and flow cytometric procedures (**Figure 5A**). Among the *FY*B*^*ES*^*/*B*^*ES*^ pbRBC sampling time points (8 samples for both *FY*B*^*ES*^*/*B*^*ES*^ #1 and *FY*B*^*ES*^*/*B*^*ES*^ #2), five samples showed rPvDBPII binding at least 2-fold above the negative control; both *FY*B*^*ES*^*/*B*^*ES*^ donors showed evidence of rPvDBPII 5-fold above the negative control. Interestingly, rPvDBPII binding was also variable for Fy-positive individuals, irrespective of their genotype. Overall, the rPvDBPII binding (fold-increase over negative control) was 32.4 for *FY*B/*B* (11.8; 110.9); 17.2 for *FY*A/*A* (4.9; 47.7); 1.9 for *FY*B*^*ES*^*/*B*^*ES*^ #1 (0.7; 5.9); 2.4 for *FY*B*^*ES*^*/*B*^*ES*^ #2 (0.5; 6.7). Of additional interest, across this series of samples from the four pbRBC donors featured, the higher levels of Fy expression on *FY*B*^*ES*^*/*B*^*ES*^ pbRBC were comparable to the lower level of rPvDBPII binding for the *FY*A/*A* pbRBC (**Figure 5A; see \***).

**Figure 5.**
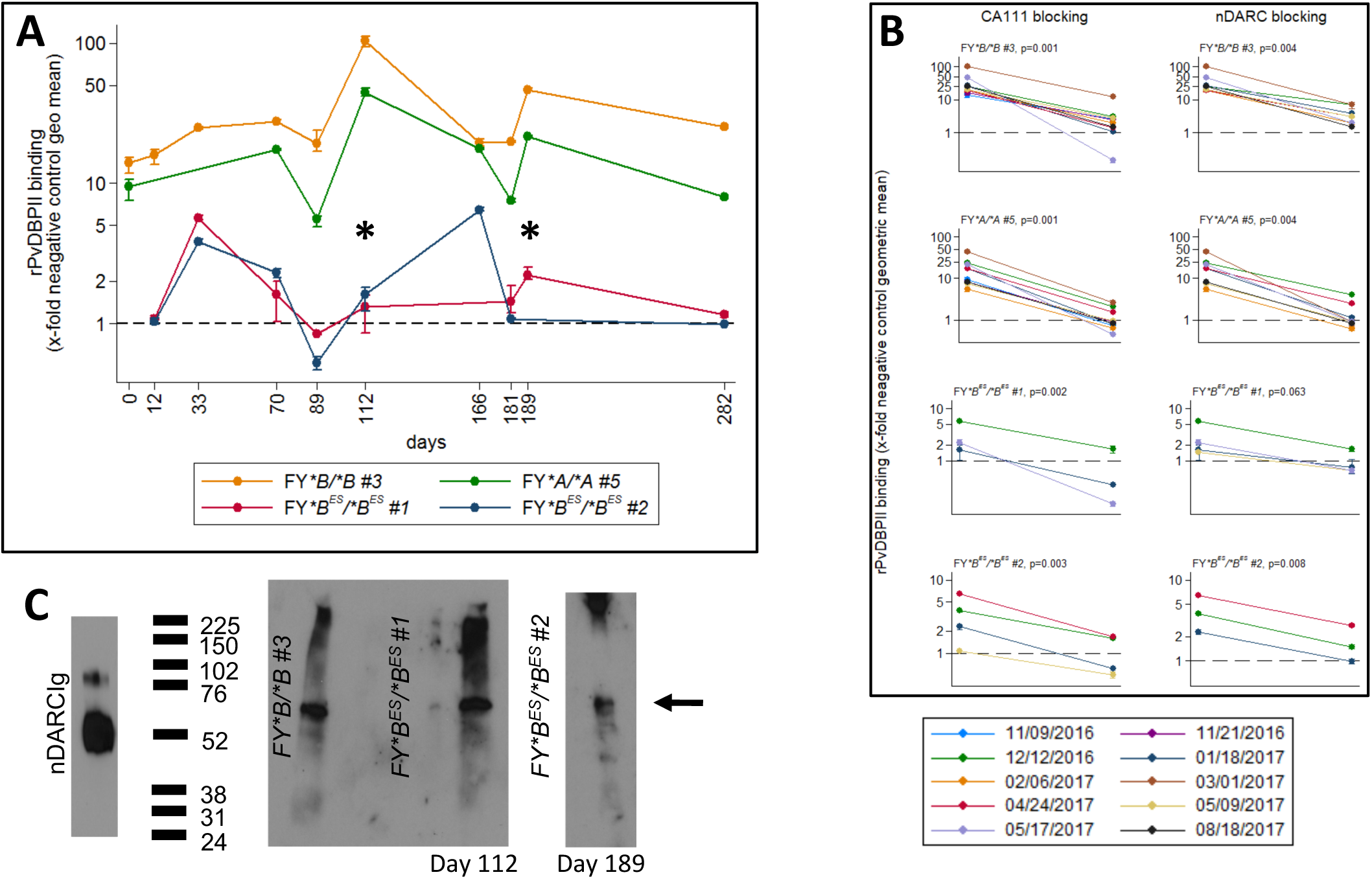
rPvDBPII binding to and Fy antigen capture from Fy-positive and Fy-negative pbRBC over time. Part A. rPvDBPII binding on Fy-positive and Fy-negative pbRBC over time. Each line corresponds to the binding of rPvDBPII on the pbRBC from one donor. Four donors were followed for up to 10 months: one *FY*B/*B*, one *FY*A/*A*, two *FY*B*^*ES*^*/*B*^*ES*^ (donors #1 and #2). At each time point, the binding was tested in triplicate and binding is represented as the x-fold increase over negative control. Each donor had its own negative control for each time point. The upper, lower and the mean values are indicated for each time point. Over time, an extensive variability of the binding has been observed for both Fy-positive and Fy-negative donors. All the experiments have been conducted using the same antibodies (same reference) and the flow cytometer (Attune NxT, Thermo Fisher). Part B. Blocking of the rPvDBPII binding to Fy-positive and Fy-negative pbRBC. Studies evaluating rPvDBPII binding to donor pbRBC were analyzed following the same protocols described for erythrocyte binding assays. The experiments performed here evaluate inhibition of rPvDBPII binding to donor pbRBC by CA111 and nDARCIg. Left Panel – Results demonstrate the inhibition of rPvDBPII binding with the addition of the Fy-specific VHH, CA111 (30ug/mL). Right Panel – Results demonstrate the inhibition of rPvDBPII binding with the addition of the Fy-IgG protein, nDARCIg (60ug/mL). Statistical analysis of this composite experiment included multiple time points for each individual sample donor: n=9 for *FY*B/*B*, n=7 for *FY*A/*A*, n=7 for *FY*B*^*ES*^*/*B*^*ES*^ #1, n=7 for *FY*B*^*ES*^*/*B*^*ES*^ #2. A paired, non-parametric Wilcoxon Signed-Ranks Test was performed to assess the differences of rPvDBPII binding without (data points on the left of each graph) or with (data points on the right of each graph) blocking (*FY*B/*B* – CA111: Z = 2.803, nDARCIg: Z = 2.521; *FY*A/*A* – CA111: Z = 2.803, nDARCIg: Z = 2.521; *FY*B*^*ES*^*/*B*^*ES*^ #1 – CA111: Z = 2.666, nDARCIg: Z = 2.197; *FY*B*^*ES*^*/*B*^*ES*^ #2 – CA111: Z = 2.380, nDARCIg: Z = 2.366). The dashed line represents the level of binding for the negative controls. Individual samples for each time point had their own negative control. Part C. Specific capture and detection of the Fy protein from the surface of pbRBC. This figure presents direct evidence of the specific capture, immunoprecipitation and detection of the Fy protein from the surface of the pbRBC of *FY*B/*B* and *FY*B*^*ES*^*/*B*^*ES*^ donors. In the left panel, nDARC is the recombinant Duffy protein. In this lane, the protein and its dimer were distinguishable. In the center panel we demonstrate CA111-based capture of the Fy protein from pbRBC of both *FY*B/*B* and *FY*B*^*ES*^*/*B*^*ES*^ (Donor #1; March 1, 2017 collection). In the right panel we demonstrate CA111-based capture of the Fy protein from pbRBC of *FY*B*^*ES*^*/*B*^*ES*^ (Donor #2; May 17, 2017 collection). The dates referred to dates for which flow cytometry results are provided Figure 4 and Figure 5. Fy protein was immunoprecipitated from pbRBC protein extract obtained (12.5×10^9^ pbRBC from both Fy-positive and Fy-negative; 2.5 mL of leukocyte depleted whole blood); leukocyte depletion confirmed by light microscopy. Detection of the CA111-captured Fy protein on these Western blots was performed using a polyclonal anti-ACKR1.

To confirm the specificity of rPvDBPII binding to pbRBC, we used CA111 to bind available Fy protein on the pbRBC and thereby block rPvDBPII access to the RBC. In a second set of experiments we used the Fy recombinant protein fused to the Fc region of IgG1, nDARCIg(39), to block rPvDBPII binding on the pbRBC surface (**Figure 5B**). For these experiments, we tested one *FY*B/*B*, one *FY*A/*A* and two *FY*B*^*ES*^*/*B*^*ES*^ donors (#1 and #2) at multiple time points. For the Fy-positive donors, the Fy-specific CA111 nanobody (Figure 5B left panel) decreased rPvDBPII binding from 25.2-fold (5.8; 110.9) over negative controls to 2.1-fold (0.2; 12.1) (P-value =0.001 for *FY*B/*B*; P-value =0.001 for *FY*A/*A*). Under these same conditions, rPvDBPII binding decreased for the *FY*B*^*ES*^*/*B*^*ES*^ pbRBC from 2.0-fold (0.5;6.4) over negative controls to 0.8-fold (0.2;1.9) (P-value =0.002 for *FY*B*^*ES*^*/*B*^*ES*^ #1; P-value =0.003 for *FY*B*^*ES*^*/*B*^*ES*^ #2).

In partner experiments to assess the specificity of rPvDBPII binding to pbRBC (**Figure 5B right panel**), nDARCIg was observed to decrease rPvDBPII binding from 25.2-fold (5.8; 110.9) over negative controls to 2.81-fold (0.5;7.7) (P-value=0.004 for *FY*B/*B*; P-value=0.004 for *FY*A/*A*). For the *FY*B*^*ES*^*/*B*^*ES*^ pbRBC, nDARC decreased rPvDBPII binding to pbRBC from 2.0-fold (0.5;6.4) over negative controls to 1.0-fold (0.3;2.8). Thus, both CA111 and nDARCIg blocked binding of PvDBPII to *FY*B*^*ES*^*/*B*^*ES*^ pbRBCs, providing evidence of Fy-specific binding of PvDBPII to *FY*B*^*ES*^*/*B*^*ES*^ pbRBC.

As further evidence of Fy protein expression on the *FY*B*^*ES*^*/*B*^*ES*^ pbRBC surface, we solubilized pbRBC membranes and immunoprecipitated the Fy protein using CA111. Results show that following CA111-capture, we successfully detected the Fy protein on Western blots probed with a second Fy-specific antibody (polyclonal anti-ACKR1) from one *FY*B/*B* and both *FY*B*^*ES*^*/*B*^*ES*^ donors (**Figure 5C**).

### Interrogating Fy protein expression on erythroid progenitor cells *in vitro*

Given a wide range of results pointing to *P. vivax* preference for infection of reticulocytes (40-48) and the apparent low, variable expression Fy protein on *Fy*B*^*ES*^*/*B*^*ES*^ pbRBC, we postulated that this expression pattern may reflect the lower and declining stages of Fy expression on aging pbRBC. Further recent evidence from murine studies showing that the Fy protein is expressed at higher levels on pro-erythroblasts and normoblasts, compared to mature RBC (49), suggesting that we should investigated Fy expression in erythroid precursors. We initiated these *in vitro* studies by expanding CD34^pos^ (expressed on megakaryocyte/erythrocyte progenitors (MEP (50-54))) from the peripheral blood of one previous *FY*B/*B* donor and *FY*B*^*ES*^*/*B*^*ES*^ Donors #1 and #2. At Days 11, 15 and 18, cells were collected to monitor erythroid lineage maturation using the surface markers CD36, CD71, Glycophorin A and Band 3 and query Fy protein expression (**Figure 6A** and **Figure S3A**). As expected, during the differentiation process, CD36 and CD71 were decreasing, whereas Glycophorin A and Band 3 increased among the most mature RBCs. No difference in the maturation stages from the different donors was observed over time (**Figure S3B**). Expression of the Fy protein during these differentiation time points was monitored by either flow cytometry (CA111 binding) and/or Western blot (CA111 immunoprecipitation; anti-ACKR1 detection). For the *FY*B/*B* donor, at Day11, half of the cell population expressed the Fy protein at the cell surface whereas after Day15 the whole population expressed the Fy protein. From the two different *FY*B*^*ES*^*/*B*^*ES*^ donors, the expression of the Fy protein was observed *in vitro* as early as Day11 of maturation. Consistent with our previous results on pbRBC, the Fy protein expression was lower on *FY*B*^*ES*^*/*B*^*ES*^ cells (11 and 35%) than on *FY*B/*B* cells (49%) (**Figure 6A**). However, we observed that significantly higher proportions of the erythroid precursor cells from *FY*B*^*ES*^*/*B*^*ES*^ donors tested positive for the Fy protein (Day18 proportion of Fy-positive cells; 49.7% for *FY*B*^*ES*^*/*B*^*ES*^ #1 and 83.8% for *FY*B*^*ES*^*/*B*^*ES*^ #2) than was observed previously for pbRBC (highest proportion of Fy-positive pbRBC from Figure 5A - 6% for *FY*B*^*ES*^*/*B*^*ES*^ #1 and 7% for *FY*B*^*ES*^*/*B*^*ES*^ #2). Expression of the Fy protein on erythroid precursors differentiated *in vitro* at Day18 was further confirmed by immunoprecipitation and Western blot (**Figure 6B and 6C**).

**Figure 6.**
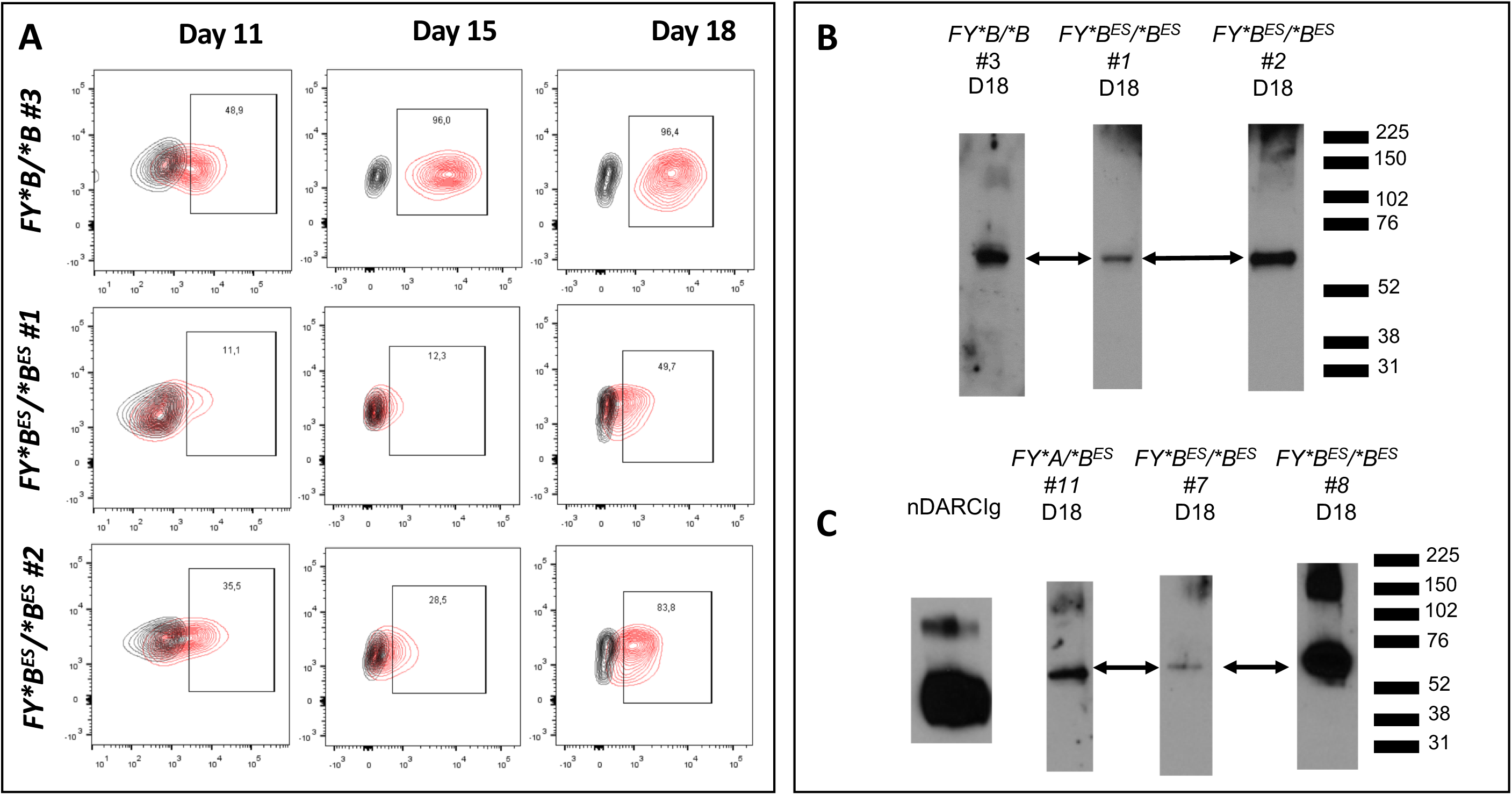
Monitoring Fy protein expression during CD34 cell *in vitro* expansion to generate erythroid precursors. Part A. CD34 positive cells were isolated from peripheral blood of one *FY*B/*B* and 2 *FY*B*^*ES*^*/*B*^*ES*^ donors (#1 and #2). CD34 positive cells isolated by immune-magnetic bead capture, were cultured and expanded for over 21 days. Cells were harvested on days 11, 15 and 18 to measure the CA111 binding during *in vitro* differentiation. For each time point and each donor, the percentage of positive cells for the binding of CA111 is expressed (population in red within the CA111-PE positive gate) in comparison to its negative control (population in black, e.g. secondary antibody only). Parts B and C. Expression of Fy protein in *FY*B*^*ES*^*/*B*^*ES*^ erythroid precursors during differentiation *in vitro* detected by Western Blot. Erythroid differentiation was initiated from CD34 positive cells isolated from peripheral blood (B) or bone marrow (C). Cells were harvested on Day18 and Fy protein was immunoprecipitated (2.0×10^5^ erythroid progenitors from both Fy-positive and Fy-negative); leukocyte depletion was not necessary as protocols for expansion of CD34 cells deplete peripheral blood leukocytes. The proteins captured by CA111 were revealed with a polyclonal anti-ACKR1 antibody (arrows). All the antibodies used to reveal the Fy protein signal were different between flow cytometry and Western blot.

These studies confirm the ability of individuals with *FY*B*^*ES*^*/*B*^*ES*^ genotype to clearly express the Fy protein in erythroid progenitor cells cultured *in vitro*; these results required confirmation of Fy expression in erythroid progenitor cells *ex vivo*.

### *Ex vivo* Fy protein expression on bone marrow erythroid precursors

The erythroid developmental stages observed during *in vitro* CD34 cell expansion studies are most commonly found in the bone marrow *in vivo*. Therefore, to evaluate Fy expression in human bone marrow, we monitored expression *ex vivo* of the following cell surface markers as accepted indicators erythroid precursors: CD34 (marker of MEP cells (50-54)), CD45 (marker to differentiate erythroid from lymphoid development (50)), CD105 (marker human erythrocyte progenitor (hEP) (54-56)), and CD71 (marker of reticulocytes (57)) cell surface markers (**Figures S4 and S5A and S5B**). To confirm observations regarding Fy expression across the developing erythroid lineage expanded from CD34 cells, we measured Fy protein expression on 10 *ex vivo* bone marrow samples (six Fy-positive and four Fy-negative). We distinguished the composition of cell lineages in each bone marrow sample, focusing on erythroid precursors and simultaneously measured the binding of CA111 by flow cytometry (**Figure 7A**). We observed CA111 recognition of the Fy protein in bone marrow erythroid precursors from both Fy-positive and Fy-negative individuals. The highest levels of CA111 binding was observed within the CD34^neg^/CD45^neg^/CD105^pos^/CD71^pos^/Band3^pos^ subpopulation (skewed toward reticulocytes; CD105 declining) for both Fy-positive and Fy-negative donors. Detailed comparative analyses of bone marrow samples from individuals representing specific *FY* genotypes are provided in **Figures S5A and S5B**. Results with CA111 have been replicated using the murine monoclonal antibody, 2C3 (**Figure 7B**). The anti-Fya and anti-Fyb antibodies used in clinical laboratories have shown limited antigen-specific binding to bone marrow. Finally, we confirmed expression of Fy protein by CA111-capture and anti-ACKR1-detection from both Fy-negative and Fy-positive donors in the erythroid precursor bone marrow population (**Figure 7C**).

**Figure 7.**
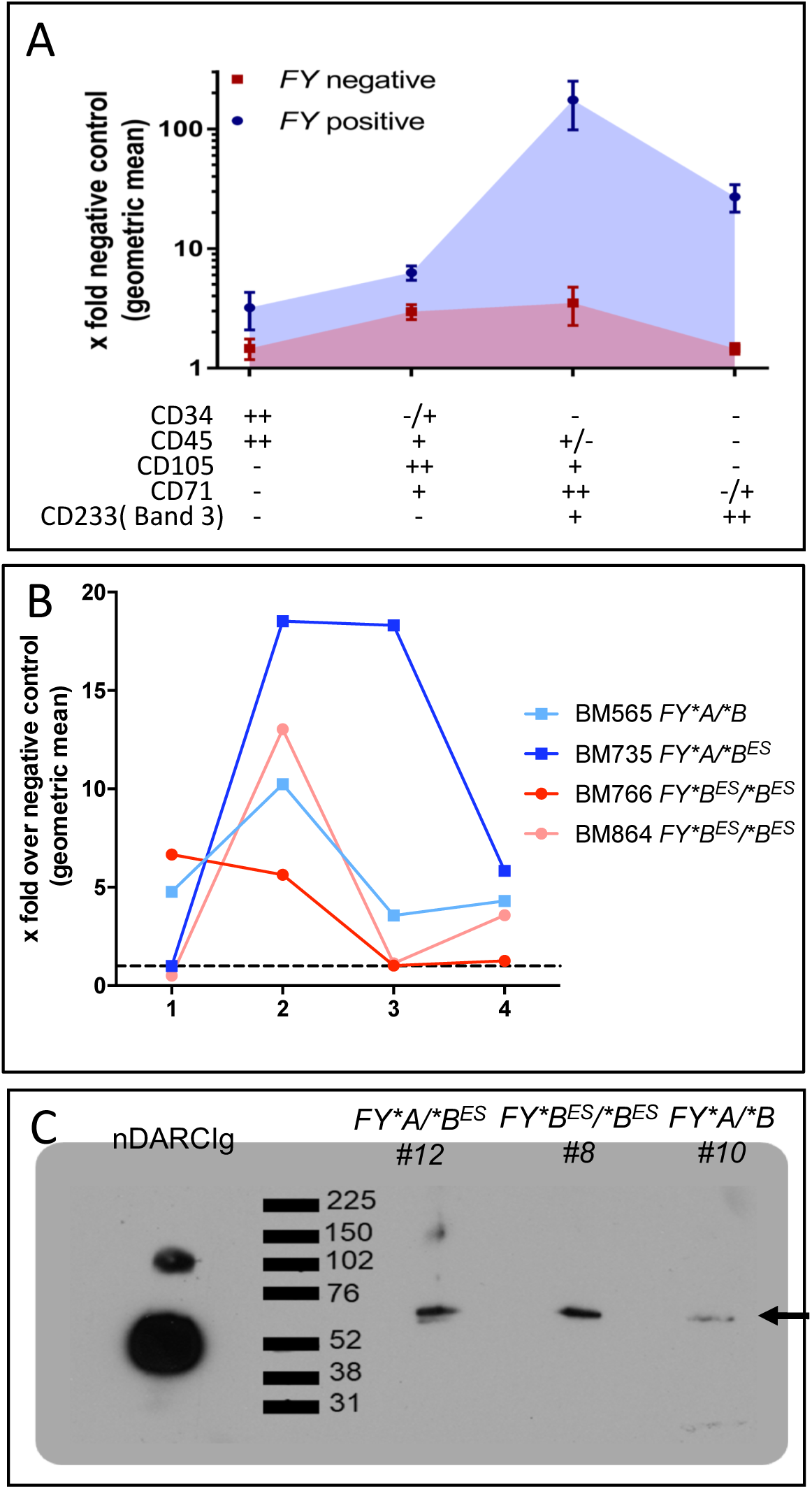
Expression of Fy protein among erythroid precursor sub-populations from bone marrow. Panel A. Four Fy-negative (in red) and six Fy-positive (in blue) bone marrows. In the bone marrow, all stages of erythroid precursors are present except mature erythrocytes (that are circulating in the peripheral blood). To determine whether a particular erythroid developmental stage expressed the Fy protein (and monitor relative expression levels), we added a panel of 5 extra-cellular markers known to distinguish bone marrow lineages. The x-axis represents the signal of those markers defining the erythroid population. The Fy protein was detected by CA111 binding. For each point, the fold-increase over negative control of the geometric mean fluorescence intensity is represented with its standard deviation. Panel B. Two Fy-negative (in red [BM766 and BM864]) and two Fy-positive (in blue [BM565 and BM735]) bone marrows. Similar to Panel A, we added a panel of 5 extra-cellular markers known to distinguish bone marrow lineages. The x-axis represents the signal of those markers defining the erythroid population. The Fy protein was detected by 2C3 binding. For each point, the fold-increase over negative control of the geometric mean fluorescence intensity is shown. Note that BM735 is a heterozygous carrier of the Fy-negative allele (*FY*A/*B*^*ES*^); BM565 is heterozygous for both Fy-positive alleles (*FY*A/*B*). Panel C. Expression of Fy protein in *Fy*B*^*ES*^*/*B*^*ES*^ CD45 negative erythroid precursors from *ex vivo* bone marrow. CD45 negative cells (erythroid precursors in bone marrow) of three bone marrow samples were sorted by flow cytometry. The membrane proteins of the sorted cells were then immunoprecipitated with the CA111 and revealed by Western Blot with a polyclonal anti-ACKR1 antibody (arrow). Both Fy-positive and Fy-negative show a signal for the Fy protein in erythroid precursors. Fy protein was immunoprecipitated from bone marrow shown to be CD45^neg^ (2.0×10^5^ erythroid progenitors from both Fy-positive and Fy-negative); leukocyte depletion was not. On the same samples, the immunoprecipitated proteins were probed with an anti-CXCR2 (see Supplemental Figure 2B) and did not show any signal, meaning that CXCR2 is not expressed on CD45 negative erythroid precursors from bone marrow.

## Discussion

Given our findings on *P. vivax* infection of Fy-negative people from Madagascar (58) and other malarious sites across Africa (25), we wanted to investigate how parasite and host invasion proteins interacted with Fy-negative RBCs. To initiate these studies, it was necessary to change how we have performed erythrocyte binding assays (34). Given that Fy-negatives are infected with *P. vivax*, we cannot assume that Fy-specific antibodies or rPvDBPII do not bind to Fy-negative RBC (22, 33, 59, 60). Additionally, because we have observed examples of varying Fy expression (King and Zimmerman; unpublished data), it was necessary to conduct a time series study to document fluctuation of Fy expression on both Fy-positive and Fy-negative RBC. This approach revealed novel findings from the outset of our studies, that the Fy protein is expressed on genotypically Fy-negative erythroid cells.

Our initial studies focused on peripheral blood RBCs (pbRBC), showed that Fy expression on Fy-negative pbRBC was periodically positive and that these RBC could interact with rPvDBPII at levels similar to RBC from *FY*A/*A* people. Well aware that numerous studies point to *P. vivax* preference for infection of reticulocytes (40, 42-48), we were interested to study Fy expression across a broader time frame of erythroid development. Recent evidence from murine and human studies shows higher levels of Fy protein expression on pro-erythroblasts and normoblasts, compared to mature RBC (46, 49). Additionally, Malleret et al. have demonstrated that immature reticulocytes (CD71^pos^), most commonly found in the bone marrow are preferentially invaded by *P. vivax* compared to older reticulocytes and erythrocytes (CD71^neg^), in the peripheral blood (47). Obaldia et al. emphasize that the bone marrow is an important reservoir of *P. vivax* infection of *Aotus* and *Saimiri* monkeys (61).

Our first studies on earlier erythroid stages focused on enriching CD34^pos^ MEPs that circulate in the peripheral blood, allowing them to mature *in vitro* under conditions favoring erythroid development(51). We purposefully included the same Fy-positive and Fy-negative people from our work demonstrating pbRBC CA111 and rPvDBPII binding to provide continuity across our study design. We felt it was particularly important to continue study of the same Fy-negative donors so that we could bridge observations from early erythroid development to mature erythrocytes. Interestingly, results showed that significantly higher proportions of the erythroid precursor cells from *FY*B*^*ES*^*/*B*^*ES*^ donors were Fy protein-positive than their pbRBC.

With observation of Fy protein expression from the CD34 cell maturation experiments, the logical *in vivo* extension of this investigation was to repeat our studies on bone marrow samples as the site for natural erythroid cell development. After identification of a strategy to segregate the earliest multipotent stem cells (CD34^pos^) differentiating toward erythroid (CD105^pos^) and not lymphoid (CD105^neg^) development, we continued to monitor expression of CD71 and Band 3 as markers distinguishing reticulocyte maturation. Results from these bone marrow studies showed Fy protein expression on early stage cells committing to erythroid development in both Fy-positive and Fy-negative people. Our observations suggest that Fy expression initiates in an early erythroid subpopulation (CD34^pos^/CD45^pos^/CD105^pos^/CD71^neg^/Band3^neg^) in both Fy-positive and Fy-negative people. This may suggest that Fy expression is driven by GATA-2, prior to proerythroblast differentiation and transition to GATA-1 (55, 62). The observed decline of Fy signal in CD34^neg^/CD45^neg^/CD105^neg^/CD71^pos^/Band3^pos^ cells, appears to be consistent with RBC extrusion of its nucleus, after which RBC precursors are no longer able to express their genes and the cell can only be equipped with the gene products expressed prior this important developmental event. With compromised *FY* gene expression associated with the GATA-1, *FY*^*ES*^ SNP, the amount of Fy protein begins to decay early and from a lower surface concentration from Fy-negative than Fy-positive erythroid cells. Our results suggest Fy6-specific CA111 is more sensitive than antibodies used in conventional serological typing.

Potential that bone marrow could be the source for reticulocytes for *in vitro* cell culture is becoming an increasingly important area of *P. vivax* research. Fernandez-Becerra et al. have expanded CD34^pos^ hematopoietic stem cells to generate CD71^pos^ reticulocytes for *in vitro P. vivax* culture (63). Others have concentrated their efforts on enriching reticulocytes from cord blood. Optimizing these *in vitro* culture systems will require additional trials as the importance of adult versus fetal hemoglobin, reticulocyte cell structure complexity and Duffy blood group polymorphism are addressed (64, 65).

Because our studies in Madagascar suggested that *P. vivax* had developed an Fy-independent RBC invasion mechanism (58), we were also interested to see that rPvDBPII binds to the surface of the different erythroid target cells used in our studies (pbRBC; expanded CD34 cell populations; bone marrow). We were further interested in the outcomes of our competitive inhibition studies. These experiments showed that addition of both CA111 and nDARCIg significantly reduced rPvDBPII binding to Fy-positive pbRBC and essentially blocked rPvDBPII binding to Fy-negative pbRBC. These latter results suggest that rPvDBPII binding may be limited to the Fy protein. If there were additional, equally attractive erythrocyte membrane proteins with which to interact, it is difficult to conclude how CA111 and nDARCIg inhibition of rPvDBPII binding could have been as complete.

### Conclusion

Our findings present evidence that the *FY* GATA-1 promoter SNP does not fully abolish erythroid lineage Fy protein expression. With increasing evidence that Fy-negative people are not completely resistant to *P. vivax* blood stage malaria (25), new reservoirs of *P. vivax* within infected individuals must be considered (47, 61). Beyond gaining new insight on RBC invasion, it is important to understand the mechanisms of pathogenesis and clinical implications of *P. vivax* malaria in the bone marrow. With extensive involvement of the bone marrow, it becomes easier to understand how significant anemia can result from *P. vivax* malaria. At the very least, peripheral blood infection may significantly underestimate the total body parasitemia of a *P. vivax* infection (66). Finally, to gain a full appreciation of the global burden of vivax malaria (67, 68), further study is required to understand how the parasite uses the poorly understood reservoir of Fy-negative people to propagate and generate gametocytes that contribute to on-going transmission.

## Methods

### Study Participants and Sample collection

Study protocols were approved by the University Hospitals of Cleveland Institutional Review Board (#08-03-33 and #09-90-195). We included eleven Fy-positive (three females and eight males) and sixteen Fy-negative persons (nine females and seven males) (**Table S1**); all Fy-negative persons were homozygous for the negative GATA-1 mutation (rs2814778; *FY*B*^*ES*^*/*B*^*ES*^). All blood and bone marrow samples were processed within 2 hours of collection.

### DNA Extraction and PCR-based Genotyping

All study participants were genotyped by previously described methods (58) (**See Supplemental Methods; Text S2**).

### Protein expression

*CA111* – Variable domain of a heavy chain of the heavy-chain only (VHH), nanobody of camelids. This nanobody was kindly provided by Olivier Bertrand (National Institute of Blood Transfusion (INTS), Paris, France) (35) (See SI Methods).

#### rPvDBPII recombinant protein (Sal I variant(69))

Region II of Salvador I strain PvDBP (rPvDBPII; 37.6 KDa) was expressed as inclusion bodies in *Escherichia coli* and then refolded and purified as previously described (70-72). The protein purified before refolding served as a negative control.

#### Duffy Antigen Receptor for Chemokine chimeric protein construct (nDARCIg Fy^b^)

Briefly, as previously described (39), HEK293H cells were co-transfected with pCDM8-DARC-Fc (first N-terminal 60 amino acids of Fy) and pRc/CMV-TPST (human sulfotransferase).

### Erythrocyte binding assays

Erythrocyte binding assays were performed based on previously developed strategies and reagents (34, 60); details provided in Supplemental Methods. In our studies, we also compare CA111 results with the murine MoMab, NaM185-2C3 (also known as 2C3; anti-Fy6) (73, 74) anti-FyA (ref: 808 186 BioRad) and anti-FyB (ref: 808 191 BioRad) antibodies.

Multiple flow cytometry strategies were followed according to specific applications in this study. Experiments evaluated CA111 and rPvDBPII binding to pbRBC in a total of 100,000 cells (all samples analyzed in triplicate). Erythrocyte binding assays were optimized by adjusting the blocking concentrations for BSA (**Figure S1A**) and through the use of unfolded rPvDBPII as a negative control (**Figure S1B**). Additionally, our assay development also tested Fy protein expression on *FY* transfected/non-transfected K562 cells (75) (expressing/not expressing, respectively [**Figure S1C**]). Following experiments on the maturation of CD34+ hematopoietic stem cells, flow cytometry evaluated at least 50,000 cells (see **Figure S3**). In evaluation of bone marrow, at least 500,000 cells were evaluated to ensure sufficient data capture for erythroid precursor subpopulations of interest.

### rPvDBPII Erythrocyte Capture Assay

This assay was previously described(76). Briefly, ∼10^6^ cells were collected from packed RBC and 10 µg of recombinant PvDBPII was added. After 2 hours of incubation at room temperature, the preparation was layered on top of 500 µL of dibutylphtalate and centrifuged at 11,000 rpm for 30 sec. The pellet was collected and the proteins bound at the RBC surface were eluted adding 20 µL of 1.5 M NaCl drop wise and shaking. After incubation (5 min), this procedure was repeated with 1.0 M of NaCl and then 0.3 M of NaCl. The preparation was centrifuged at 14,000 rpm for 2 min. The supernatant was collected and the proteins were separated on a Tris-Glycine SDS 4-20% gel (Biorad) in reducing conditions and then transferred onto a PVDF membrane. The membrane was blocked for one hour with TBS 0.1% tween, 5% milk, incubated with rabbit anti-PvDBPII (1: 2,000), washed, then followed by the secondary antibody anti-rabbit HRP (Thermo Scientific). Chemiluminescent signal was detected using SuperSignal™ (Pierce) on X-ray film.

### Erythroid differentiation of CD34 positive cells

Isolation of CD34 positive cells was performed from peripheral blood donors (*FY*B/*B* and *FY*B*^*ES*^*/*B*^*ES*^ donors; #1 and #2) and from bone marrow samples. Mononuclear cells were isolated by Ficoll Plaque Plus (GE Healthcare) and frozen in 10%DMSO. The thawed cells were positively selected for CD34 by magnetic beads (human CD34 MicroBead Kit; Miltenyi Biotech). Between 10,000 and 100,000 cells were obtained after selection and differentiated in culture over 21 days. A detailed protocol is published by Jingping Hu *et al* (51). The *in vitro* differentiation was monitored by flow cytometry with 400,000 cells per day (Day 11, Day 15, Day 18) monitoring surface markers CD36 (BD Biosciences), CD71 (BD Biosciences), Band 3 (American Research Product) and Glycophorin A (Life Technology).

### Bone marrow cell preparation and staining

Fresh bone marrow was collected in heparin. Erythroid precursors were isolated by Ficoll Paque Plus. A panel constituted by CD45 PerCP Cy5.5, CD71 APC Cy7, CD34 FITC, Band3 PE and CD105 APC markers were used to differentiate erythroid precursors in bone marrow.

### Erythroid cell protein extraction and Western blot analysis

Extracting the Fy protein from the surface of erythroid cells was conducted to specifically capture evidence of *FY* gene expression. These experiments were performed on pbRBCs from Fy-positive and Fy-negative donors. Target cell populations included pbRBC, expanded CD34 cell populations and bone marrow. Details of protein extraction, immunoprecipitation and Western blot analyses are provided in Supplemental Methods.

### Statistical Methods

Statistical analyses were performed using STATA version 13.0

## Acknowledgments

We are grateful to the study donors who patiently participated over the course of this investigation and to David N. Wald, Director of the Hematopoietic Biorepository & Cellular Therapy Core and Basabi Maitra for recruitment of bone marrow donors. Howard Meyerson, Keith E. Shults and the Comprehensive Cancer Center of Case Western Reserve University and University Hospitals of Cleveland generously contributed to the flow cytometry study design and provided technical support. We thank Chetan Chitnis for providing nDARCIg; Olivier S. Bertrand and Sylvie Cochet (INSERM/University Paris Diderot) for the gift of anti-Fy antibodies (2C3 and CA111) and objective criticisms integral to this study; Anne-Lyse Baudrier, Emmanuel Collec and the Centre National de Reference pour les Groupes Sanguins (CNRGS) for providing the anti-FyA and anti-FyB antibodies. We finally thank Allison M. Shackelford for graphic art assistance with the visual abstract overview of our manuscript.

This study was funded by grants from the Veterans Affairs Research Service (BX001350) and NIH Grant (R01 AI064687) to CLK and an NIH Grant (R01 AI097366) to PAZ.

The authors have declared that no conflict of interest exists.

## Author contributions

CD, SD, BG, CLK and PAZ conceived the study; CD and SD wrote the first draft of the paper with PAZ. CD, SD, CLK, PAZ and JWJ developed the flow cytometry design for the bone marrow analysis. SB conducted all genotyping assays. SK and RF conducted erythrocyte binding assays and flow cytometry. LC conducted Western blot analyses. AG produced unfolded rPvDBPII. NHT, EC and NDS synthesized rPvDBPII. YC developed and provided 2C3 and CA111. SM, SD and BG conducted in vitro proliferation, expansion, and differentiation of CD34+ cells. CLK and PAZ oversaw all aspects of the study. CD, SD, BG, CLK and PAZ analyzed the results. All authors read and approved the final manuscript.

## REFERENCES

1. Donahue RP, Bias WB, Remwick JH, & McKusick VA (1968) Probable assignment of the Duffy blood group locus to chromosome 1 in man. Proc Natl Acad Sci U S A 61:949–955.

2. Bachelerie F, et al. (2014) International Union of Basic and Clinical Pharmacology. [corrected]. LXXXIX. Update on the extended family of chemokine receptors and introducing a new nomenclature for atypical chemokine receptors. Pharmacol Rev 66(1):1–79.

3. Darbonne WC, et al. (1991) Red blood cells are a sink for interleukin 8, a leukocyte chemotaxin. J Clin Invest 88(4):1362–1369.

4. Dawson TC, et al. (2000) Exaggerated response to endotoxin in mice lacking the Duffy antigen/receptor for chemokines (DARC). Blood 96(5):1681–1684.

5. Neote K, Darbonne W, Ogez J, Horuk R, & Schall TJ (1993) Identification of a promiscuous inflammatory peptide receptor on the surface of red blood cells. J Biol Chem 268(17):12247–12249.

6. Reutershan J, Harry B, Chang D, Bagby GJ, & Ley K (2009) DARC on RBC limits lung injury by balancing compartmental distribution of CXC chemokines. Eur J Immunol 39(6):1597–1607.

7. Vergara C, et al. (2008) Gene encoding Duffy antigen/receptor for chemokines is associated with asthma and IgE in three populations. American journal of respiratory and critical care medicine 178(10):1017–1022.

8. Hub E & Rot A (1998) Binding of RANTES, MCP-1, MCP-3, and MIP-1alpha to cells in human skin. Am J Pathol 152(3):749–757.

9. Middleton J, et al. (1997) Transcytosis and surface presentation of IL-8 by venular endothelial cells. Cell 91(3):385–395.

10. Pruenster M, et al. (2009) The Duffy antigen receptor for chemokines transports chemokines and supports their promigratory activity. Nat Immunol 10(1):101–108.

11. Rot A (1992) Endothelial cell binding of NAP-1/IL-8: role in neutrophil emigration. Immunol Today 13(8):291–294.

12. Zarbock A, et al. (2007) The Duffy antigen receptor for chemokines in acute renal failure: A facilitator of renal chemokine presentation. Crit Care Med 35(9):2156–2163.

13. O’Leary PA (1927) Treatment of neurosyphilis by malaria: Report on the three years’ observation of the first one hundred patients treated. Journal of the American Medical Association 89(2):95–100.

14. Boyd MF & Stratman-Thomas WK (1933) Studies on benign tertian malaria. 4. On the refractoriness of negroes to inoculation with Plasmodium vivax. Am J Hyg 18:485–489.

15. Young MD, Ellis JM, & Stubbs TH (1946) Studies on imported malarias. 5. Transmission of foreign Plasmodium vivax by Anopheles quadrimaculatus. Am J Trop Med 26:477–482.

16. Young MD, Eyles DE, Burgess RW, & Jeffery GM (1955) Experimental testing of the immunity of negroes to Plasmodium vivax. J. Parasitol 41:315–318.

17. Bray RS (1957) Studies on Plasmodium ovale in Liberia. American Journal of Tropical Medicine and Hygiene 6:961–970.

18. Bray RS (1958) The susceptibility of Liberians to the Madagascar strain of Plasmodium vivax. J. Parasitol 44:371–373.

19. Miller LH, Mason SJ, Clyde DF, & McGinniss MH (1976) The resistance factor to Plasmodium vivax in blacks. The Duffy-blood-group genotype, FyFy. N Engl J Med 295(6):302–304.

20. Tournamille C, Colin Y, Cartron JP, & Le Van Kim C (1995) Disruption of a GATA motif in the Duffy gene promoter abolishes erythroid gene expression in Duffy-negative individuals. Nat Genet 10(2):224–228.

21. Zimmerman PA, et al. (1999) Emergence of FY*A(null) in a Plasmodium vivax-endemic region of Papua New Guinea. Proc Natl Acad Sci U S A 96(24):13973–13977.

22. Chitnis CE & Miller LH (1994) Identification of the erythrocyte binding domains of Plasmodium vivax and Plasmodium knowlesi proteins involved in erythrocyte invasion. J Exp Med 180(2):497–506.

23. Ranjan A & Chitnis CE (1999) Mapping regions containing binding residues within functional domains of Plasmodium vivax and Plasmodium knowlesi erythrocyte-binding proteins. Proc Natl Acad Sci U S A 96(24):14067–14072.

24. Aikawa M, Miller LH, Johnson J, & Rabbege J (1978) Erythrocyte entry by malarial parasites. A moving junction between erythrocyte and parasite. J Cell Biol 77(1):72–82.

25. Zimmerman PA (2017) Plasmodium vivax Infection in Duffy-Negative People in Africa. Am J Trop Med Hyg 97(3):636–638.

26. Woolley IJ, Hotmire KA, Sramkoski RM, Zimmerman PA, & Kazura JW (2000) Differential expression of the duffy antigen receptor for chemokines according to RBC age and FY genotype. Transfusion 40(8):949–953.

27. Lopez GH, et al. (2015) A novel FY*A allele with the 265T and 298A SNPs formerly associated exclusively with the FY*B allele and weak Fy(b) antigen expression: implication for genotyping interpretative algorithms. Vox Sang 108(1):52–57.

28. Olsson ML, et al. (1998) The Fy(x) phenotype is associated with a missense mutation in the Fy(b) allele predicting Arg89Cys in the Duffy glycoprotein. Br J Haematol 103(4):1184–1191.

29. Parasol N, et al. (1998) A novel mutation in the coding sequence of the FY*B allele of the Duffy chemokine receptor gene is associated with an altered erythrocyte phenotype. Blood 92(7):2237–2243.

30. Tournamille C, et al. (1998) Arg89Cys substitution results in very low membrane expression of the Duffy antigen/receptor for chemokines in Fy(x) individuals. Blood 92(6):2147–2156.

31. Mallinson G, Soo KS, Schall TJ, Pisacka M, & Anstee DJ (1995) Mutations in the erythrocyte chemokine receptor (Duffy) gene: the molecular basis of the Fya/Fyb antigens and identification of a deletion in the Duffy gene of an apparently healthy individual with the Fy(a-b-) phenotype. Br J Haematol 90(4):823–829.

32. Rios M, et al. (2000) New genotypes in Fy(a-b-) individuals: nonsense mutations (Trp to stop) in the coding sequence of either FY A or FY B. Br J Haematol 108(2):448–454.

33. Wertheimer SP & Barnwell JW (1989) Plasmodium vivax interaction with the human Duffy blood group glycoprotein: identification of a parasite receptor-like protein. Exp Parasitol 69(4):340–350.

34. King CL, et al. (2011) Fy(a)/Fy(b) antigen polymorphism in human erythrocyte Duffy antigen affects susceptibility to Plasmodium vivax malaria. Proc Natl Acad Sci U S A 108(50):20113–20118.

35. Smolarek D, et al. (2010) A recombinant dromedary antibody fragment (VHH or nanobody) directed against human Duffy antigen receptor for chemokines. Cell Mol Life Sci 67(19):3371–3387.

36. Horuk R (2015) The Duffy Antigen Receptor for Chemokines DARC/ACKR1. Front Immunol 6:279.

37. Haynes JD, et al. (1988) Receptor-like specificity of a Plasmodium knowlesi malarial protein that binds to Duffy antigen ligands on erythrocytes. J Exp Med 167(6):1873–1881.

38. Miller LH, Hudson D, & Haynes JD (1988) Identification of Plasmodium knowlesi erythrocyte binding proteins. Mol Biochem Parasitol 31(3):217–222.

39. Choe H, et al. (2005) Sulphated tyrosines mediate association of chemokines and Plasmodium vivax Duffy binding protein with the Duffy antigen/receptor for chemokines (DARC). Mol Microbiol 55(5):1413–1422.

40. Kitchen SF (1938) The infection of reticulocytes by Plasmodium vivax. Am J Trop Med 18:347–353.

41. Garnham PCC (1966) Malaria Parasites and Other Haemosporidia (Blackwell Scientific Publications, Oxford) p 1114.

42. Mons B (1990) Preferential invasion of malarial merozoites into young red blood cells. Blood Cells 16(2-3):299–312.

43. Russell B, et al. (2011) A reliable ex vivo invasion assay of human reticulocytes by Plasmodium vivax. Blood 118(13):e74–81.

44. Lim C, et al. (2016) Reticulocyte Preference and Stage Development of Plasmodium vivax Isolates. J Infect Dis 214(7):1081–1084.

45. Cho JS, et al. (2016) Unambiguous determination of Plasmodium vivax reticulocyte invasion by flow cytometry. Int J Parasitol 46(1):31–39.

46. Ovchynnikova E, et al. (2017) DARC extracellular domain remodeling in maturating reticulocytes explains Plasmodium vivax tropism. Blood 130(12):1441–1444.

47. Malleret B, et al. (2015) Plasmodium vivax: restricted tropism and rapid remodeling of CD71-positive reticulocytes. Blood 125(8):1314–1324.

48. Gruszczyk J, et al. (2018) Transferrin receptor 1 is a reticulocyte-specific receptor for Plasmodium vivax. Science 359(6371):48–55.

49. Duchene J, et al. (2017) Atypical chemokine receptor 1 on nucleated erythroid cells regulates hematopoiesis. Nat Immunol 18(7):753–761.

50. Kondo M, et al. (2003) Biology of hematopoietic stem cells and progenitors: implications for clinical application. Annu Rev Immunol 21:759–806.

51. Hu J, et al. (2013) Isolation and functional characterization of human erythroblasts at distinct stages: implications for understanding of normal and disordered erythropoiesis in vivo. Blood 121(16):3246–3253.

52. Attar A (2014) Changes in the Cell Surface Markers During Normal Hematopoiesis: A Guide to Cell Isolation. Global Journal of Hematology and Blood Transfusion 1:20–28.

53. Sidney LE, Branch MJ, Dunphy SE, Dua HS, & Hopkinson A (2014) Concise review: evidence for CD34 as a common marker for diverse progenitors. Stem Cells 32(6):1380–1389.

54. Mori Y, Chen JY, Pluvinage JV, Seita J, & Weissman IL (2015) Prospective isolation of human erythroid lineage-committed progenitors. Proc Natl Acad Sci U S A 112(31):9638–9643.

55. Machherndl-Spandl S, et al. (2013) Molecular pathways of early CD105-positive erythroid cells as compared with CD34-positive common precursor cells by flow cytometric cell-sorting and gene expression profiling. Blood Cancer J 3:e100.

56. Wangen JR, Eidenschink Brodersen L, Stolk TT, Wells DA, & Loken MR (2014) Assessment of normal erythropoiesis by flow cytometry: important considerations for specimen preparation. International journal of laboratory hematology 36(2):184–196.

57. Kono M, Kondo T, Takagi Y, Wada A, & Fujimoto K (2009) Morphological definition of CD71 positive reticulocytes by various staining techniques and electron microscopy compared to reticulocytes detected by an automated hematology analyzer. Clin Chim Acta 404(2):105–110.

58. Menard D, et al. (2010) Plasmodium vivax clinical malaria is commonly observed in Duffy-negative Malagasy people. Proc Natl Acad Sci U S A 107(13):5967–5971.

59. Singh S, et al. (2001) Biochemical, biophysical, and functional characterization of bacterially expressed and refolded receptor binding domain of Plasmodium vivax duffy-binding protein. J Biol Chem 276(20):17111–17116.

60. Tran TM, et al. (2005) Detection of a Plasmodium vivax erythrocyte binding protein by flow cytometry. Cytometry A 63(1):59–66.

61. Obaldia N, 3rd, et al. (2018) Bone Marrow Is a Major Parasite Reservoir in Plasmodium vivax Infection. MBio 9(3).

62. Ferreira R, Ohneda K, Yamamoto M, & Philipsen S (2005) GATA1 function, a paradigm for transcription factors in hematopoiesis. Mol Cell Biol 25(4):1215–1227.

63. Fernandez-Becerra C, et al. (2013) Red blood cells derived from peripheral blood and bone marrow CD34(+) human haematopoietic stem cells are permissive to Plasmodium parasites infection. Mem Inst Oswaldo Cruz 108(6):801–803.

64. Malleret B, et al. (2011) A rapid and robust tri-color flow cytometry assay for monitoring malaria parasite development. Scientific reports 1:118.

65. Noulin F, et al. (2012) Cryopreserved reticulocytes derived from hematopoietic stem cells can be invaded by cryopreserved Plasmodium vivax isolates. PLoS One 7(7):e40798.

66. Aitken G (1943) Sternal puncture in the diagnosis of malaria. Lancet 242(6268):466–468.

67. Howes RE, et al. (2016) Global Epidemiology of Plasmodium vivax. Am J Trop Med Hyg.j

68. Howes RE, et al. (2015) Plasmodium vivax Transmission in Africa. PLoS neglected tropical diseases 9(11):e0004222.

69. Collins WE, Contacos PG, Krotoski WA, & Howard WA (1972) Transmission of four Central American strains of Plasmodium vivax from monkey to man. J Parasitol 58(2):332–335.

70. Batchelor JD, et al. (2014) Red blood cell invasion by Plasmodium vivax: structural basis for DBP engagement of DARC. PLoS pathogens 10(1):e1003869.

71. Batchelor JD, Zahm JA, & Tolia NH (2011) Dimerization of Plasmodium vivax DBP is induced upon receptor binding and drives recognition of DARC. Nat Struct Mol Biol 18(8):908–914.

72. Chen E, et al. (2016) Broadly neutralizing epitopes in the Plasmodium vivax vaccine candidate Duffy Binding Protein. Proc Natl Acad Sci U S A 113(22):6277–6282.

73. Wasniowska K, et al. (1996) Identification of the Fy6 epitope recognized by two monoclonal antibodies in the N-terminal extracellular portion of the Duffy antigen receptor for chemokines. Mol Immunol 33(11-12):917–923.

74. Wasniowska K, et al. (2002) Structural characterization of the epitope recognized by the new anti-Fy6 monoclonal antibody NaM 185-2C3. Transfus Med 12(3):205–211.

75. Chaudhuri A, et al. (1994) Expression of the Duffy antigen in K562 cells. Evidence that it is the human erythrocyte chemokine receptor. J Biol Chem 269(11):7835–7838.

76. Wickramarachchi T, Devi YS, Mohmmed A, & Chauhan VS (2008) Identification and characterization of a novel Plasmodium falciparum merozoite apical protein involved in erythrocyte binding and invasion. PLoS One 3(3):e1732.

