## Supplemental Information for "Duffy Antigen Expression in Erythroid Bone Marrow Precursor Cells of Genotypically Duffy Negative Individuals"

**Supplemental Methods –**

1. Study Participants (Table S1)
2. Development of CA111 (Text S1)
3. Comparison of Fy-specific antibody reagents (Figure S1)
4. Erythroid cell protein extraction and Western blot analysis (Figure S2)
    a. Protein extraction
    b. Immunoprecipitation
    c. Western blot
5. Maturation and expansion of CD34+ Hematopoietic Stem Cells (Figure S3)
6. Flow cytometry of erythroid progenitor subpopulations in bone marrow (Figure S4 & S5)
7. DNA-based genotyping (Text S2)
8. Erythrocyte binding Flow Cytometry assays (Figure S6)

### 1.0 Study Participants

| Table S1. Study Donor Information | | | | | | |
|---|---|---|---|---|---|---|
| ID | Tissue | Sample | Sex | Age | Genotype | Serotype |
| #1 | Peripheral Blood | D2 | F | 21 | $FY*B^{ES}/*B^{ES}$ | na |
| #2 | Peripheral Blood | D3 | F | 21 | $FY*B^{ES}/*B^{ES}$ | na |
| #3 | Peripheral Blood | BB | M | 38 | *FY*B/*B* | Fya-b+ |
| #4 | Peripheral Blood | AB | F | 36 | *FY*A/*B* | Fya+b+ |
| #5 | Peripheral Blood | AA | M | 27 | *FY*A/*A* | na |
| #6 | Bone Marrow | BM_863 | M | 62 | $FY*B^{ES}/*B^{ES}$ | na |
| #7 | Bone Marrow | BM_864 | M | 41 | $FY*B^{ES}/*B^{ES}$ | na |
| #8 | Bone Marrow | BM_874 | M | 37 | $FY*B^{ES}/*B^{ES}$ | na |
| #9 | Bone Marrow | BM_766 | F | 39 | $FY*B^{ES}/*B^{ES}$ | na |
| #10 | Bone Marrow | BM_885 | F | 35 | *FY*A/*B* | na |
| #11 | Bone Marrow | BM_786 | M | 31 | *FY*B/*B* | na |
| #12 | Bone Marrow | BM_797 | M | 31 | $FY*A/*B^{ES}$ | na |
| #13 | Bone Marrow | BM_838 | M | 62 | $FY*A/*B^{ES}$ | na |
| #14 | Bone Marrow | BM_881 | F | 37 | $FY*A/*B^{ES}$ | na |
| #15 | Bone Marrow | BM_875 | M | 33 | $FY*A/*B^{ES}$ | na |
| #16 | Bone Marrow | BM_735 | M | 45 | $FY*A/*B^{ES}$ | na |
| #17 | Bone Marrow | BM_565 | M | 40 | *FY*A/*B* | na |
| #18 | Frozen RBC | 36253 | M | na | $FY*B^{ES}/*B^{ES}$ | Fya-b- |
| #19 | Frozen RBC | 5873 | F | na | $FY*B^{ES}/*B^{ES}$ | Fya-b- |
| #20 | Frozen RBC | 31395 | M | na | $FY*B^{ES}/*B^{ES}$ | Fya-b- |
| #21 | Frozen RBC | 20636 | M | na | $FY*B^{ES}/*B^{ES}$ | Fya-b- |
| #22 | Frozen RBC | 12725 | F | na | $FY*B^{ES}/*B^{ES}$ | Fya-b- |
| #23 | Frozen RBC | 12664 | F | na | $FY*B^{ES}/*B^{ES}$ | Fya-b- |
| #24 | Frozen RBC | 39207 | F | na | $FY*B^{ES}/*B^{ES}$ | Fya-b- |
| #25 | Frozen RBC | 6293 | M | na | $FY*B^{ES}/*B^{ES}$ | Fya-b- |
| #26 | Frozen RBC | 21667 | F | na | $FY*B^{ES}/*B^{ES}$ | Fya-b- |
| #27 | Frozen RBC | 24947 | F | na | $FY*B^{ES}/*B^{ES}$ | Fya-b- |

**Rare *FY* Genotypes** - Unsuccessful efforts were made to obtain viable erythroid cells from donors homozygous for reported rare nonsense SNPs and deletions in the *FY* gene coding region (1, 2) from three international blood centers.  These include, the New York Blood Center, New York, the National Institute of Blood Transfusion (INTS), Paris, France and the International Blood Group Reference Laboratory (International Rare Donor Panel), Bristol, UK.  Laboratory Directors indicated that while their databases include in excess of 500,000 donors, their roles either no longer included individuals with these rare genotypes or that these individuals were no longer living.

**2.0. Development of CA111 – variable domain of a heavy chain of the heavy-chain only (VHH), nanobody of camelids (Text S1).**

Smolarek et al. develop new tools for Fy studies based on the variable domain of heavy chain-only antibodies (VHH) produced by camelids, to produce a single-domain antigen-specific binding fragment (3). Camelid immunoglobulins are composed of only heavy chain segments and lack all light chains (4). The variable region of these heavy chains-only antibodies is easily cloned from naive or immunized camelid lymphocytes. Resulting fragments, known as VHHs or nanobodies, are the smallest intact antigen-binding fragment derivative endowed with the properties of an authentic antibody (5).

CA111 was obtained by subcloning the CA52 VDJ sequence into a Novagen pET 28-b derived plasmid allowing expression in the cytoplasm of SHuffle (C3029H) cells. This plasmid encodes (from 5' to 3') a polyhistidine tail, a thrombin cleavage site, the VHH, and an HA-tag; it is used for routine subcloning of other VHHs using the PstI and Eco91I sites (6)

The resulting nanobody, exhibits the following characteristics.

- Interferes with the interleukin-8 binding to Fy on RBCs.
- Interferes with *P. vivax* infection of red blood cells.
- Upon linkage to a solid substrate, this nanobody acts as a powerful adsorbent to purify native Fy in a single step from a detergent extract of cells.
- Pepscan analysis identified $^{22}$**FEDVW**$^{26}$ as the peptide sequence of the linear epitope.

A single dromedary was immunized with the N-terminal extracellular domain constructs of Fy expressed in E. coli (referred to as ECD1; Amino acids 6 [H] to 63 [P] – HRAELSPSTENSS QLDFEDVWNSSYGVNDSFPDGDY[G/D]ANLEAAAPCHSCNLLDDSALP) (Smolarek et al., Cell Mol Life Sci 67:3371-3387; 2010). This domain covers the region to which Fy6 epitope, chemokines and the *P. vivax* Duffy binding protein (PvDBP) bind (amino acids 19 to 25 – $^{19}$**QLDFEDV**$^{25}$) and carries the polymorphic G/D site at amino acid 42, responsible for the Fya/Fyb allotypes. From the immunized animal, a VHH library of dromedary lymphocytes was exposed to ECD1 to yield several Fy-specific VHHs. Smolarek et al. focused on one VHH, referred to as CA52. The linear epitope of CA52 exhibits capacity to recognize the glycosylated Fy protein present on human cells, although the immunogen was a non-glycosylated polypeptide.

#### 3.0 Comparison of Fy-specific antibody reagents.

Antibody reagents available to this study have included CA111 (described above), NaM185-2C3 clone (also known as 2C3) (7, 8) as well as Anti-FyA (ref: 808 186 BioRad) and Anti-FyB (ref: 808 191 BioRad) antibodies, kindly provided by Anne-Lyse Baudrier, Emmanuel Collec and the Centre National de Reference pour les Groupes Sanguins (CNRGS). As indicated above, Pepscan analyses show that CA111 and 2C3 recognize identical amino acid sequence ($^{22}$**FEDVW**$^{26}$) (3, 9), consistent with Fy6 epitope (anti-Fy6 (10)) appears to be no longer readily available to the research community). Details of the erythrocyte binding assays shown below are described in Section 4.2. In the results shown below, we demonstrate the comparative specificities and sensitivities of CA111 and 2C3.

In **Figure S1A**, our results show that the camelid nanobody, CA111 competitively inhibited binding of 2C3 to Fya-b+ pbRBCs in a dose-dependent fashion; similarly, 2C3 competitively inhibited binding of CA111 to these same Fya-b+ pbRBCs (data not shown). These results demonstrate the Fy6-binding specificity of CA111 and 2C3. In **Figure S1B**, our results compare the binding of CA111, 2C3, anti-Fya (BioRad 808-186) and anti-Fyb (BioRad 808 191) to pbRBCs from 10 Fy-negative (all $FY^*B^{ES}/^*B^{ES}$) and two Fy-positive (one *FY*B/*B*, one *FY*A/*B*) individuals. These results show the relative differences of CA111, 2C3, anti-Fya and anti-Fyb binding to Fy-positive and Fy-negative pbRBCs. As we have shown in figures presented in the main body of the manuscript, antibody binding to Fy-positive pbRBC is substantially greater than observed to Fy-negative pbRBCs. Additionally, among pbRBC from 10 Fy-negative ($FY^*B^{ES}/^*B^{ES}$) individuals, we observed binding of CA111 to be 2 to 3-fold greater than binding to an irrelevant camelid nanobody targeting a *P. falciparum* protein. In further experiments, we observed clear binding of 2C3, anti-Fya and anti-Fyb antibodies to pbRBCs of Fy-positive individuals, but, we observed negligible binding of these antibodies to pbRBCs of $FY^*B^{ES}/^*B^{ES}$ individuals, suggesting that CA111 is more sensitive in detecting the Fy antigen than these other antibody probes when evaluating Fy expression on pbRBCs.

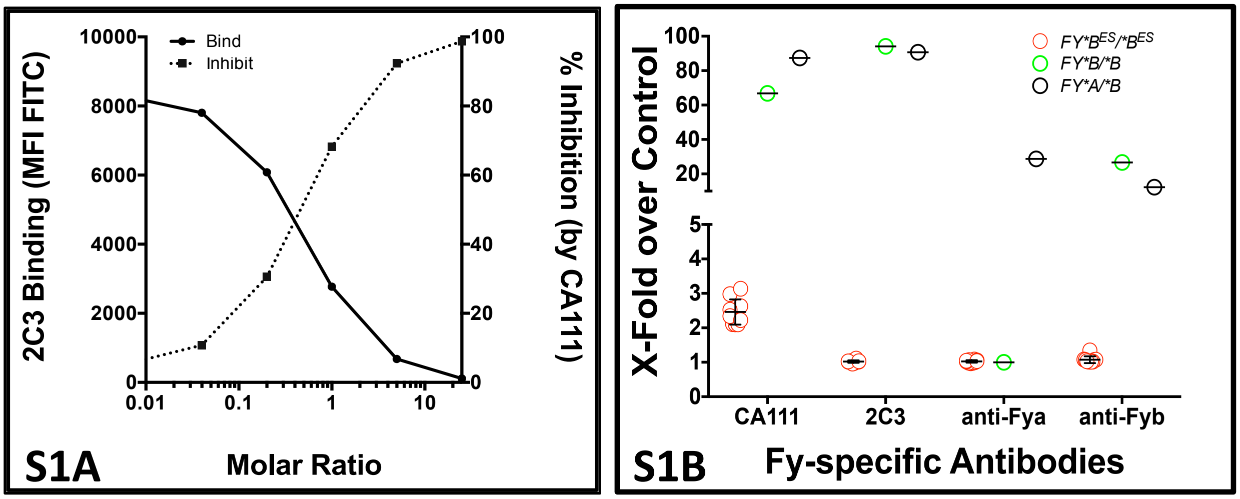


**Figure S1A. Competition assays between Fy6-specific 2C3 and CA111 binding to peripheral blood RBC (pbRBC) from Fy-positive donors.** Binding inhibition studies were performed on pbRBCs from an *FY*B/*B* donor. In this binding-inhibition study, pbRBCs were preincubated with CA111 followed by incubation with a constant concentration of 2C3 (molar ratios of CA111 to 2C3 – 0.04M to 25M). Detection of 2C3 was performed as previously described (11).

**Figure S1B. Comparison of Fy-specific antibody binding to peripheral blood RBC (pbRBC) from Fy-positive and Fy-negative donors.** The rationale for this study was to determine the pbRBC binding characteristics of CA111 in comparison to 2C3 and to Fya-specific (BioRad 808-186) and Fyb-specific (BioRad 808 191) antibodies. Blood donors available for these experiments included two Fy-positive (one *FY*A/*B, one FY*B/*B*) and ten Fy-negative (all $FY{*}B^{ES}/{*}B^{ES}$) individuals. pbRBC were prepared for flow cytometry following the optimized protocol for erythrocyte binding assays.

#### 4.0 Erythroid cell solubilization, protein extraction and Western blot analysis

*Solubilization and protein extraction* – Packed RBC (~5 mL); cells matured during the *in vitro* erythroid differentiation; and CD45 negative sorted-cells from the bone marrow were lysed by three cycles of freeze/thaw followed by sonication. After adding 5 mM Tris·HCl, pH 8.0 with protease inhibitor (Sigma Aldrich), the protein preparation was centrifuged 20 min at 16,000 × g at 4°C. The pellet was resuspended twice in 0.1 M $Na_2CO_3$ pH 11.5 with protease inhibitor and centrifuged 20 min at 16,000 × g at 4°C. To solubilized the membrane proteins from the pellet, 1mM triton X100 + protease inhibitors were added and incubated at room temperature for 30 min.

*Immunoprecipitation of Fy antigen* - CA111 was covalently coupled to magnetic beads (Dynabeads M-270 Carboxylic Acid, Invitrogen) for 30 min at room temperature with slow rotation. EDC at 100mg/ml (1-ethyl-3-[3-dimethylaminopropyl] carbodiimide) was dissolved 100mM MES (2-(N-morpholino) ethanesulfonic acid)) buffer (pH 5.0) for the activation reaction and incubated for 2 hr at 4°C with slow rotation. The solubilized extract was incubated with the activated beads for 1h. Beads were resuspended in Laemmli buffer for SDS gel and western blot or in PBS for trypsin digestion.

*Western blot* - The beads + extract solutions were separated on a Tris-Glycine SDS 4-20% gel (Biorad) in reducing conditions and were then transferred on a PVDF membrane. The membrane was blocked for one hour with TBS 0.1% Tween 5% milk. Polyclonal rabbit anti-ACKR1 (2ug/ml; LS Bioscience) and HRP anti-rabbit antibody (diluted 1/10,000 into 10 ml TBS 0.1% Tween 5% milk; Thermo Scientific) were used to probe Fy from the immunoprecipitated elution. The signal was detected on X-ray films using SuperSignal (Pierce).

*Specificity of CA111* - Tests of specificity were performed on recombinant proteins and antibodies by direct Western blotting (**Figure S2A**), or immunoprecipitating cells from patient bone marrow and subsequent Western Blot (**Figure S2B**). For the direct probe of the Western Blot, 0.1 µg of nDARCIg and rCXCR2 were loaded on a gel; 0.2 µg/mL of CA111 were used to probe the Western blot abd reveal the signal. For the immunoprecipitation experiment, 0.5 µg of rCXCR2 were used. The anti-

CXCR2 antibody (R&D system, 0.5 µg/mL) was used to probe recCXCR2 on a Western Blot as a positive control.

The resulting Western blot was probed with anti-CXCR2. Controls in the two left lanes show that anti-CXCR2 antibody did not bind to nDARCIg, but did bind to rCXCR2. To confirm that CA111 did not pull down CXCR2 from human cells, we tested the exact same samples as in Figure 7 of the main text (sorted CD45 negative cells from bone marrow samples). Results show that CA111 did not capture CXCR2 during immunoprecipitation as revealed by the absence of anti-CXCR2 signal in the three donor bone marrow samples (three right lanes

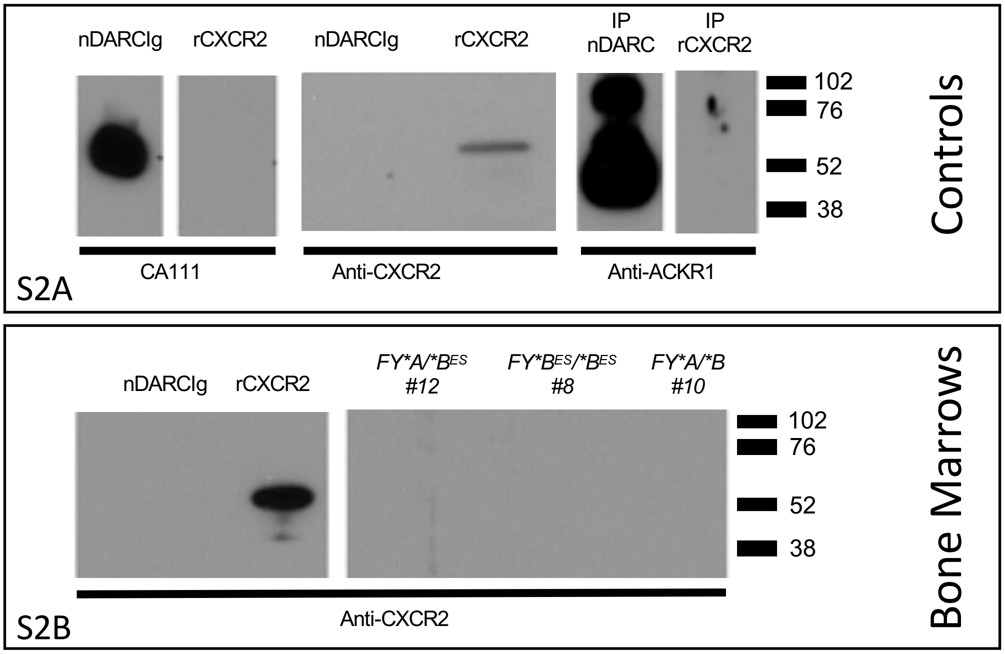


**Figure S2. Binding specificity of CA111 to recombinant proteins nDARC and rCXCR2.**

**Panel S2A.** Results show the characteristics of Fy-specific and CXCR2-specific antibody binding. CA111 binds to nDARCIg and does not bind to rCXCR2 (left). The anti-rCXCR2 antibody binds to rCXCR2 and not to nDARCIg (center). When CA111 was used to immunoprecipitate (IP) nDARCIg and rCXCR2, the resulting Western blot probed with a second Fy-specific antibody (anti-ACKR1) bound to nDARCIg and did not bind to rCXCR2 (right).

**Panel S2B.** CA111 was used to purify native Fy in a single step from a detergent extract of bone marrow cells from donors characterized by different *FY* genotypes. The resulting Western blot was probed with anti-CXCR2. Controls in the two left lanes show that anti-CXCR2 antibody did not bind to nDARC, but did bind to rCXCR2. To confirm that CA111 does not pull down CXCR2 from human cells, we have tested the exact same samples as in Figure 7 of the main text (sorted CD45 negative cells from bone marrow samples). Results show that CA111 did not capture CXCR2 during immunoprecipitation as revealed by the absence of anti-CXCR2 signal in the three donor bone marrow samples (three right lanes); results include bone marrow from both Fy-positive and Fy-negative people.

**5.0 Monitoring maturation and expansion of CD34+ Hematopoietic Stem Cells (HSCs).**

Figure S3A. Expansion of CD34 cell *in vitro*

| Day in Culture | *FY*B $^{ES}$/*B $^{ES}$* #1 | *FY*B $^{ES}$/*B $^{ES}$* #2 | *FY*B/*B* #3 |
|---|---|---|---|
| 0 | $0.1x10^6$ | $0.1x10^6$ | $0.1x10^6$ |
| 11 | $13x10^6$ | $1.8x10^6$ | $24x10^6$ |
| 15 | $20x10^6$ | $9.5x10^6$ | $32.5x10^6$ |
| 18 | $35x10^6$ | $13x10^6$ | $56x10^6$ |

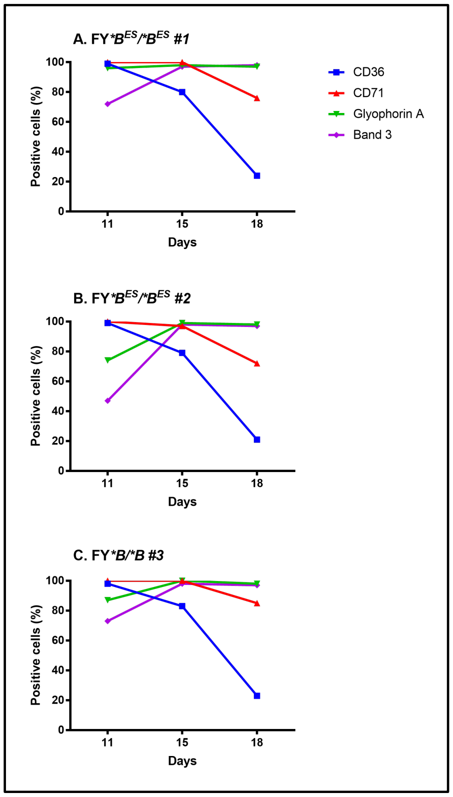


**Figure S3A** provides data on the maturation of CD34+ HSCs. Mononuclear cells were isolated by Ficoll Plaque Plus (GE Healthcare) and frozen in 10%DMSO. The thawed cells were positively selected for CD34 by magnetic beads (human CD34 MicroBead Kit; Miltenyi Biotech). Between 10,000 and 100,000 cells were obtained after selection and differentiated in culture over 21 days. A detailed protocol is published by Jingping Hu *et al*. (12) The *in vitro* differentiation was monitored by flow cytometry with 400,000 cells per day (Day 11, Day 15, Day 18) monitoring surface markers CD36 (BD Biosciences), CD71 (BD Biosciences), Band 3 (American Research Product) and Glycophorin A (Life Technology).

**Figure S3B. Monitoring of CD34 positive cells differentiation *in vitro*.**

CD34+ erythroid precursors cells were collected from the peripheral blood of 3 donors (1 Fy-positive and 2 Fy-negatives). The cells were differentiated *in vitro* the differentiation was monitored by the presence/absence of the surface markers CD36, CD71, Glycophorin A and Band 3. As expected, during the differentiation process, the cells are losing the signal CD36 and CD71 whereas

Glycophorin A and Band 3 are increasing. Overall, the cells from all the donors were observed to be at comparable stages over time. For the *FY*B/*B* donor and 2 *FY*B$^{ES}$/*B$^{ES}$* donors, the cells were counted by light microscopy during the *in vitro* differentiation at different time points (Day 0, 11, 15 and 18).

**6.0 Flow cytometric analysis of erythroid progenitor subpopulations in bone marrow.**

Erythroid precursors from fresh bone marrow samples were isolated by Ficoll Paque Plus (GE Healthcare) separation. Cells were first blocked for 30 min with FcR blocking reagent (Miltenyi – 20 µL per sample) following manufacturer's recommendations. rPvDBPII and CA111 binding experiments were performed as described above. Sub-populations of erythroid precursors were defined using directly-labeled antibodies - anti-CD45 PerCP Cy5.5 (Becton Dickinson), anti-CD34 FITC, anti-CD105 APC, anti-CD71 APC Cy7 (all from Biolegend) and an anti-Band3 PE (American Research Product). Data were acquired with a BD Biosciences LSR II (San Jose, CA) or a Thermo Fisher Attune Nxt (Waltham, MA) operated within manufacturer's specifications, which was tested with performance beads. To ensure sufficient relevant data, we aimed to acquire 500,000 cell events. Data were analyzed with FlowJo 10.0 (Treestar, Ashland, OR) or WinList 9.0 (Verity Software House, Topsham, ME). Two types of analysis were performed. In the first, used for the data presented in Figures 1-7 (main text) and Figures S1-S5, and S10, forward scatter (FSC-A) and side scatter (SSC-A) plots were used to select erythroid populations then standard gating was done to select positive and negative population from orthogonal quadrants on bivariate plots or univariate histograms. For each of the samples analyzed, unstained cells were used to evaluate cell auto-fluorescence. Unstained and single antibody stained cells, or unstained or single stained antibody-binding beads, were used to set fluorescence compensation. Negative controls for CD234 (Fy) were "secondary only" with either CA111 or rPvDBPII omitted in otherwise complete assays. Secondary and tertiary reagents were PE labeled anti HA-tag for CA111and PE labeled goat anti-rabbit, respectively for rPvDBPII (with a polyclonal anti-PvDBP secondary) . Results collected following comparisons with these negative controls allowed for calculating fold-increase of CA111or rPvDBPII over the negative control or calculating the fraction of positive cells.

In the second analysis, performed on bone marrow samples, we analyzed a subset of bone marrow donors to show the level of expression of CD34, CD45, CD105, CD71, and Fy (CD234). The antibodies and staining protocol was as above.  However, the sub-populations were gated multi-dimensionally using a series of bivariate plots to create a sequential series of erythroid differentiation states.  The fluorescence value of each parameter was divided by a correlated measure of cell size (forward scatter) providing a measure of antigen density.  The expression level of Fy was made more accurate by subtracting the fluorescence of the negative control on a subpopulation basis (median fluorescence – median background fluorescence).  The primary gating strategy is shown in **Figure S4**.

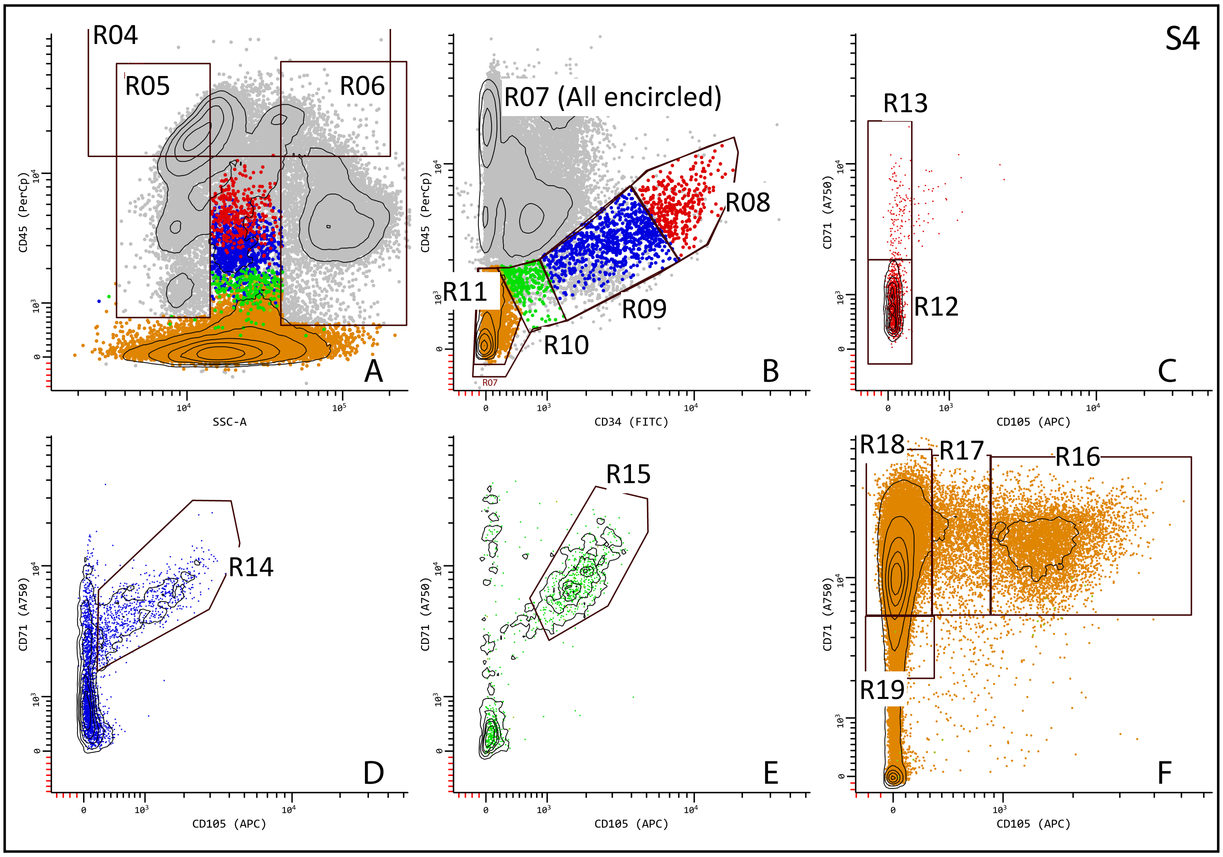


The gating strategy follows the logic presented by Machherndal-Spandl et al. (13), using ergodic principles (14) and manual data classification that mimics probability state modeling (15) and unsupervised seriation techniques (16).  The idea that dynamic expression can be derived from static snapshots is well supported (17, 18).  First, single cell events are included with a primary gate on FSC-A vs FSC-H (not shown).  Then, as shown in **Figure S4A**, cells with lymphoid or mature myeloid granularity and CD45 expression are excluded (R04, R05, R06) with Boolean NOT gates.  Next, shown in **Figure S4B**, myeloid and erythroid precursors are included and subdivided as a function of decreasing CD34 and CD45 intensity (R08, R09, R10, R11; R07 is an all-inclusive gate (R08, R09, R10, R11) used for monitoring but not data analysis). The CD34+/CD45+ precursors are further restricted based on the beginning expression of CD71 and CD105 (R12, R13, R14, R15, **Figure S4C, D, E**). Finally, erythroid cells are defined by the correlated levels of CD105 and CD71 (R16, R17. R18, F19, **Figure 4F**). From these regions, 8 sequential differentiation/maturation states are defined by Boolean logic as follows:

State 1: R08 & R12 &! (R04 | R05 | R06)

State 2: R08 & R13 &! (R04 | R05 | R06)

State 3: R09 & R14 &! (R04 | R05 | R06)

State 4: R10 & R15 &! (R04 | R05 | R06)

State 5: R07 & R16 &! (R04 | R05 | R06)

State 6: R07 & R17 &! (R04 | R05 | R06)

State 7: R07 & R18 &! (R04 | R05 | R06)

State 8: R07 & R19 &! (R04 | R05 | R06)

The results are shown in **Figures S5A and S5B (following page)**. The primary analysis as presented in Figure 7 of the main text was presented similar to this secondary analysis. Figure 7 of the main text is more conventional and statistically more robust but less informative. The goal of this secondary analysis was to create expression profiles to determine the consistency of expression dynamics between donors or different genotypes and to determine the overall differentiation-dependent expression of Fy on bone marrow precursors and generate a “target” value representing the combined expression level and population size in the bone marrow.

Gray = FSC; Green = CD34, Purple = CD45, Magenta = CD105, Auburn = CD71, Orange = Fy; Fy-negative donors (- - - - - -).

The levels of CD34, CD45, CD105, CD71, and Fy antigen (CA111 reactivity) expression density were calculated by dividing fluorescence intensity by FSC-A intensity on a single cell basis. These median values for each immune- phenotype state are shown normalized and as a function of state in **Figure S5A**. This provides an overall view of the population expression dynamics, an idea of the uniformity of the classification scheme, and points to a possible abnormality in Fy expression dynamics in Fy-negative donors.

A more biologically correct view is shown in **Figure S5B**, which takes advantage of the proportionality of the detection frequency and time in immunophenotype state. This view is not exact in that cell division is not accounted for, but it has the advantage that the fraction of cells resident in the bone marrow at expression density “X” can be discerned from the plot. Further, by integrating the expression density data (not normalized), the “target size “ expression of Duffy antigen can be reduced to a single value representing both expression dynamics and expression levels between donors (**Figure S5B**, bar plot). This analysis suggests that Duffy-negative donors express from 1-4% of the average target value of WT donors.

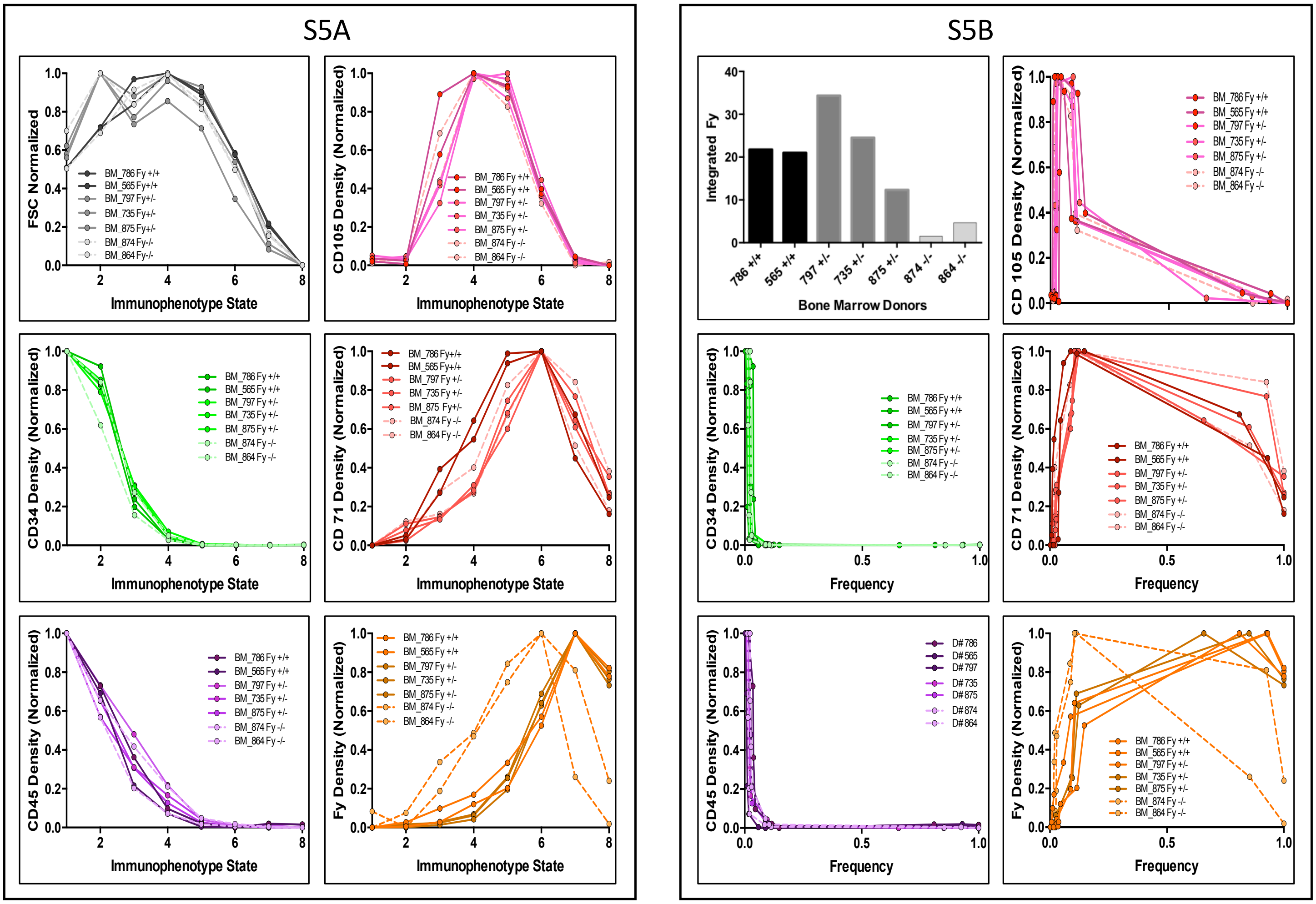


These data support and emphasize the conclusion of text Figure 7, that Fy antigen is expressed at low levels in $FY*B^{ES}/*B^{ES}$ bone marrow precursor cells and at significantly higher levels in cells from donors with at least one wild type allele. We have been able to measure Fy positivity even in a donor with very low expression (1% of average target value). Further, tentatively, Fy antigen appears to be aberrantly regulated with the peak levels occurring at an earlier stage than either wild type genotype supporting the possibility that the bone marrow is a niche of *P. vivax* infection. None of the expression dynamics can be attributed to size, which is uniform for all three genotypes and as expected for the erythroid differentiation/maturation series.

**7.0. DNA-based Genotyping (Text S2).**

Genomic DNA was extracted from the whole blood samples using the QIAGEN QIAmp DNA Blood Kit following recommended protocols with a starting blood volume of 200 µL and an elution volume of 200 µL of Buffer AE (Valencia, CA).

PCR amplifications of Duffy blood group gene sequences were performed in reaction mixtures (28µL) with 3 µL of genomic DNA, 180 µM each dNTP, 67mM Tris-HCl (pH 8.8), 6.7 mMMgSO4, 16.6mM$(NH_4)_2$SO4, 10mM 2-mercaptoethanol, 0.1 µM each primer (Primers inclusive of promoter (rs2814778) and *FY*A*/*FY*B* (rs12075) snps - forward primer, Duffy-200up 5′-CAGGCAGTGGGCGTGGG-3′; reverse primer, Duffy +730dn 5′-CTGCTAGCTAGGATACCCAG-3′; Primers inclusive of *FY*B*/*FY*X* (rs34599082) snp - forward primer, FYBXup 5′-AGCACTGTCCTCTTCATGCTTT-3′; reverse primer, FYBXdn 5′-GCAGAGCTGCGAGTGCTAC-3′) and 2.5 units of thermostable DNA polymerase under the following conditions: 95 °C for 2min, followed by 40 cycles of 95 °C (30 s), 60 °C (30 s), and 72 °C (90 s) and a final extension at 72 °C (5 min) (PCR products 912 and 1,033 bp). Following PCR amplification, products were further processed by a ligation detection reaction (LDR). The LDR was performed in a reaction mixture (15 µL) containing 20mMTris-HCl buffer (pH 7.6), 25 mM potassium acetate, 10mM magnesium acetate, 1mM NAD+, 10mM DTT, 0.1% Triton X-100, 13nM each LDR probe,1 µL of PCR product, and 2 units of Taq DNA ligase (New England BioLabs). LDR probes consisted of six allele-specific oligonucleotides and three fluorescently labeled conserved-sequence oligonucleotides. The allele-specific probes contained an MTAG sequence for further hybridization with complementary sequence oligonucleotides bound to Luminex FlexMAP fluorescent microspheres; conserved-sequence probes were 5′ phosphorylated and 3′ biotinylated.

*Sequences of the LDR probes used were as follows:*

Duffy null promoter snp (rs2814778) at nucleotide −67 (t, wild-type; c, erythrocyte silent):

PRO ntT MTAG_A018: 5′-ACACTTATCTTTCAATTCAATTACcattagtccttggctcttat-3′

PRO ntC MTAG_A020: 5′- CTTTCTCATACTTTCAACTAATTTtcattagtccttggctcttac-3′

PRO common: 5′-phosphate-cttggaagcacaggcgctg-biotin-3′.

Codon 42 snp (rs12075) associated with Fya and Fyb phenotypes (g, Fya; a, Fyb):

*FY*A* ntG MTAG-A022: 5′- CAAACAAACATTCAAATATCAATCttcccagatggagactatgg-3′

*FY*B* ntA MTAG-A026: 5′- TACATTCAACACTCTTAAATCAAActtcccagatggagactatga-3′

Codon42 common: 5′-phosphate-tgccaacctggaagca-biotin-3′.

Codon 89 snp (rs34599082) associated with Fyb or the Fyb$^{weak}$ phenotypes (c, Fyb; t, Fyb$^{weak}$):

*FY*B* ntC MTAG-A028: 5′- CACTTAATTCATTCTAAATCTATCtgcttttcagacctctctcc-3′

*FY*X* ntT MTAG-A034: 5′- ACTTATTTCTTCACTACTATATCAtgcttttcagacctctctct-3′

Codon89 common: 5′-phosphate-gctggcagctctgccctggct-biotin-3′.

**8.0 Erythrocyte binding assays** - Erythrocyte binding assays were performed based on previously developed strategies and reagents (19, 20).

Peripheral blood RBC ($1x10^6$) were first blocked for 30 min with PBS 2% BSA and then further incubated with **CA111** (30 µg/mL final concentration; 20 min at 37°C); this nanobody includes a human influenza hemagglutinin (HA) epitope tag. These pbRBC were washed two times with PBS 0.2% BSA and then incubated with mouse anti-HA tag (Biolegend; 30 µL of a 1:1000 dilution in PBS 0.2% BSA; 20 min at 37°C). The pbRBC were washed again (2x with PBS 0.2% BSA). Finally, pbRBC were incubated with a goat, anti-mouse phycoerythrin (PE)-tagged antibody (eBioscience; 40 µl of a 1:80 dilution in PBS 0.2% BSA; 20 min at 37°C in the dark). Following 2 washes in PBS 0.2% BSA and a final wash in PBS, cells were analyzed by flow cytometry. For *ex vivo* fresh bone marrow samples, a direct anti-HA tag phycoerythrin (PE)-tagged antibody was used to reduce the background (Miltenyi, dilution 1:11). Similar procedures were followed for other Fy-specific antibody reagents.

Peripheral blood RBC ($1x10^6$) were first blocked for 30 min with PBS 2% BSA and then further incubated with **PvDBPII Sal1** (20µg/mL final concentration; 60 µL of a 1:20 dilution in PBS 0.2% BSA; 30 min at 37°C). After 2X washes with PBS 0.2%BSA, these cells were incubated with a rabbit polyclonal anti-PvDBP serum (30 µL of a 1:50 dilution in PBS 0.2% BSA; 20 min at 37°C). Cells were washed again 2X with PBS 0.2%BSA and then incubated with rabbit phycoerythrin (PE)-tagged antibody (Sigma; 40 µL of a 1:32 dilution in PBS 0.2% BSA; before 20 min at 37°C in the dark). After 2 washes in PBS 0.2% BSA and a final wash in PBS, cells were subjected to analytical flow cytometry. Erythrocyte binding assays were further optimized by adjusting the blocking concentrations for BSA (**Figure S6A**) and through the use of unfolded rPvDBPII as a negative control (**Figure S6B**).

Finally, to test detection of Fy by CA111 on cells that do express, against cells that do not express this protein, we compared *FY* transfected/non-transfected K562 cells (21) (respectively (**Figure S6C**).

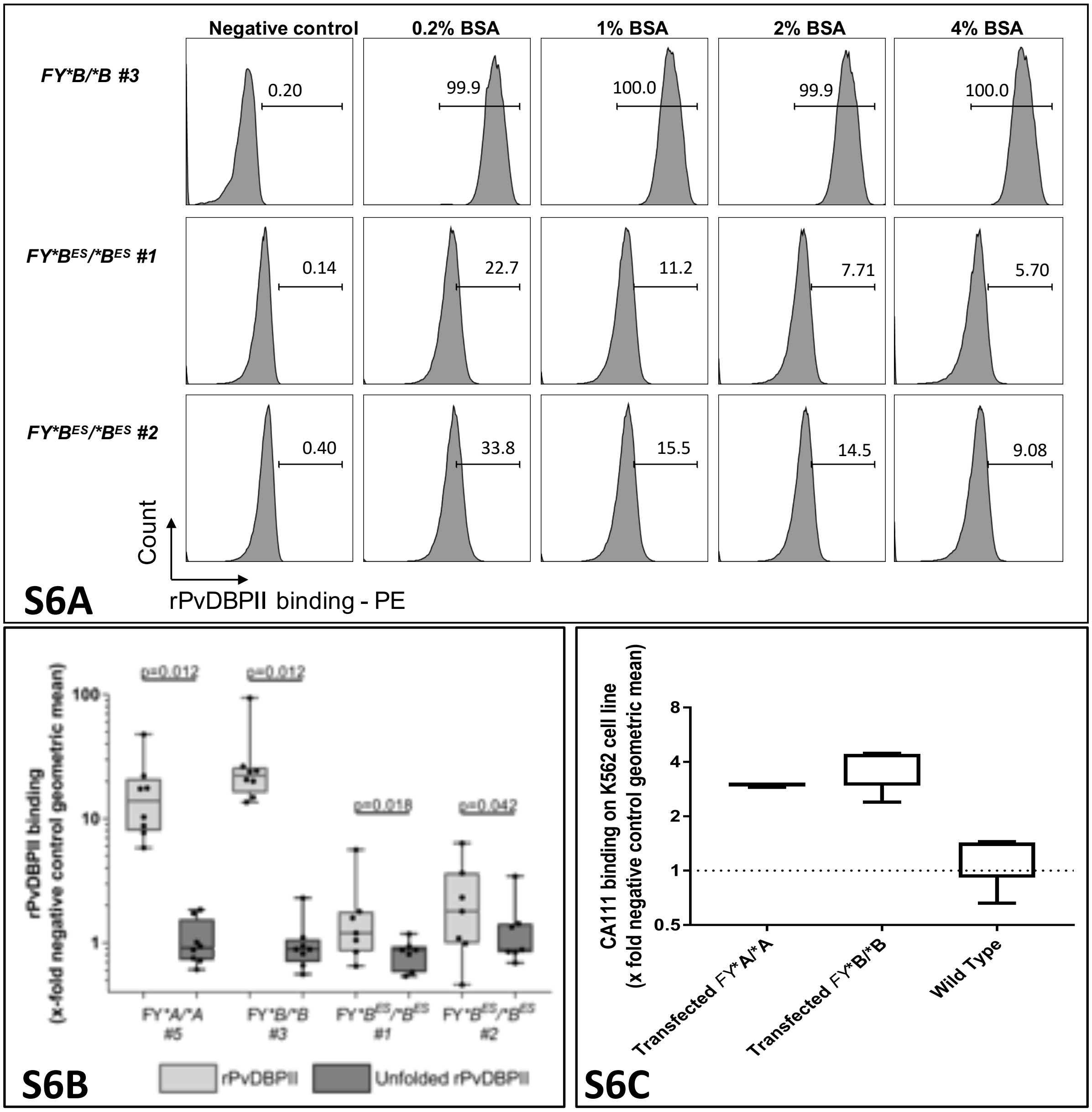


**Figure S6A. The effect of increasing BSA concentration on rPvDBPII binding to peripheral blood RBC (pbRBC) from Fy-positive and Fy-negative donors.** The rationale for this study was to determine if it was possible to reduce or eliminate non-specific rPvDBPII binding to donor pbRBC with increasing BSA concentrations during pre-incubation steps in erythrocyte binding assays; particular interest was focused on the results for *FY*B^ES/*B^ES* (donors #1 and #2). The binding of rPvDBPII to pbRBC was monitored by flow cytometry by comparing rPvDBPII-specific fluorescence of negative controls (minus rPvDBPII; plus secondary anti-rPvDBPII, rabbit polyclonal antibody; plus anti-rabbit phycoerythrin-conjugateted goat antibody) to rPvDBPII-specific fluorescence at increasing BSA concentrations (0.2%1.0%, 2.0% and 4.0%). The percentage of positive cells for each individual pbRBC donor is indicated above the gate determined for positive rPvDBPII binding. Results show that while rPvDBPII binding was reduced approximately 2-fold between 0.2% to 1.0% BSA for *FY*B$^{ES}$/*B$^{ES}$* donors (statistic), rPvDBPII-specific fluorescence was not further reduced (statistic) nor eliminated from the pbRBC for *FY*BES/*BES* (donors #1 and #2). With these results in mind, the BSA pre-incubation blocking concentration for flow cytometry of erythrocyte binding assays was performed using 1.0% BSA for studies focused on pbRBC and 2.0% for bone marrow.

**Figure S6B. Comparison of rPvDBPII binding between folded and unfolded protein in *FY*A/*A* or *FY*B/*B* and *FY*B$^{ES}$/*B$^{ES}$* red blood cells.** To evaluate if the observed rPvDBPII binding was statistically different than controls, we compared the binding of folded rPvDBPII to the binding of the unfolded protein (same amino acid construct) on RBC. We found that the binding observed in all Duffy genotypes was higher for refolded rPvDBPII compared to the unfolded rPvDBPII. This figure represents rPvDBPII binding (light gray boxes) or unfolded rPvDBPII binding (dark gray boxes) on 4 different individuals; specific individual data points appear as fill black circles. Non-parametric paired sign tests were performed to test the differences between the medians per individual (rPvDBPII folded vs unfolded). The number of measures was n=8 for each of the Fy-positive individuals and n=7 for each of the 2 Fy-negative donors. Wilcoxon matched pairs sign rank test. *FY*A/*A*: z= 2.521 ; *FY*B/*B*: z= 2.521 ; *FY*B$^{ES}$/*B$^{ES}$* #1: z= 2.366 ; *FY*B$^{ES}$/*B$^{ES}$* #2: z= 2.028. Comparisons among unfolded rPvDBPII interaction among all four study participants showed no significant difference (non-parametric median test Pearson Chi²: p=0.853).

rPvDBPII binding inhibition assays on peripheral red blood cells (pbRBC) were also performed in the context of chymotrypsin treatment. For these experiments, erythrocytes were pre-treated with chymotrypsin (1 mg/mL) 1 h at 37°C. The enzyme treatment was stopped by washing 5 times with PBS 0.2%BSA. Treated and untreated red cells were blocked in PBS 1% BSA for 30 min at 37°C and washed 2 times with PBS 0.2% BSA.

**Figure S6C. Comparison of CA111 to transfected and non-transfected K562 cells.** Results show the fold over negative control binding of CA111 to human erythroleukemia K562 line transfected (K562-*FY*A* and K562-*FY*B*) and non-transfected cells (Wild Type). Cell surface expression of Duffy was detected in transfected cells; however, no Duffy protein was detected in non-transfected cells. These control experiments illustrate flow cytometry outcomes when cells are not capable of expressing a Duffy protein on their cell surface shown by the non-transfected cells.

### References


1. Mallinson G, Soo KS, Schall TJ, Pisacka M, & Anstee DJ (1995) Mutations in the erythrocyte chemokine receptor (Duffy) gene: the molecular basis of the Fya/Fyb antigens and identification of a deletion in the Duffy gene of an apparently healthy individual with the Fy(a-b-) phenotype. *Br J Haematol* 90(4):823-829.
2. Rios M, *et al.* (2000) New genotypes in Fy(a-b-) individuals: nonsense mutations (Trp to stop) in the coding sequence of either FY A or FY B. *Br J Haematol* 108(2):448-454.
3. Smolarek D, *et al.* (2010) A recombinant dromedary antibody fragment (VHH or nanobody) directed against human Duffy antigen receptor for chemokines. *Cell Mol Life Sci* 67(19):3371-3387.
4. Hamers-Casterman C, *et al.* (1993) Naturally occurring antibodies devoid of light chains. *Nature* 363(6428):446-448.
5. Saerens D, *et al.* (2005) Engineering camel single-domain antibodies and immobilization chemistry for human prostate-specific antigen sensing. *Anal Chem* 77(23):7547-7555.
6. Habib I, *et al.* (2013) V(H)H (nanobody) directed against human glycophorin A: a tool for autologous red cell agglutination assays. *Anal Biochem* 438(1):82-89.
7. Wasniowska K, *et al.* (1996) Identification of the Fy6 epitope recognized by two monoclonal antibodies in the N-terminal extracellular portion of the Duffy antigen receptor for chemokines. *Mol Immunol* 33(11-12):917-923.
8. Wasniowska K, *et al.* (2002) Structural characterization of the epitope recognized by the new anti-Fy6 monoclonal antibody NaM 185-2C3. *Transfus Med* 12(3):205-211.
9. Tournamille C, *et al.* (2003) Structure-function analysis of the extracellular domains of the Duffy antigen/receptor for chemokines: characterization of antibody and chemokine binding sites. *Br J Haematol* 122(6):1014-1023.
10. Nichols ME, Rubinstein P, Barnwell J, Rodriguez de Cordoba S, & Rosenfield RE (1987) A new human Duffy blood group specificity defined by a murine monoclonal antibody. Immunogenetics and association with susceptibility to Plasmodium vivax. *J Exp Med* 166(3):776-785.
11. Menard D, *et al.* (2010) Plasmodium vivax clinical malaria is commonly observed in Duffy-negative Malagasy people. *Proc Natl Acad Sci U S A* 107(13):5967-5971.

12. Hu J, *et al.* (2013) Isolation and functional characterization of human erythroblasts at distinct stages: implications for understanding of normal and disordered erythropoiesis in vivo. *Blood* 121(16):3246-3253.

13. Machherndl-Spandl S, *et al.* (2013) Molecular pathways of early CD105-positive erythroid cells as compared with CD34-positive common precursor cells by flow cytometric cell-sorting and gene expression profiling. *Blood Cancer J* 3:e100.

14. Wheeler RJ (2015) Analyzing the dynamics of cell cycle processes from fixed samples through ergodic principles. *Mol Biol Cell* 26(22):3898-3903.

15. Bagwell CB, *et al.* (2015) Human B-cell and progenitor stages as determined by probability state modeling of multidimensional cytometry data. *Cytometry B Clin Cytom* 88(4):214-226.

16. Saeys Y, Van Gassen S, & Lambrecht BN (2016) Computational flow cytometry: helping to make sense of high-dimensional immunology data. *Nat Rev Immunol* 16(7):449-462.

17. Avva J, Weis MC, Sramkoski RM, Sreenath SN, & Jacobberger JW (2012) Dynamic expression profiles from static cytometry data: component fitting and conversion to relative, "same scale" values. *PLoS ONE* 7(7):e38275.

18. Jacobberger JW, Avva J, Sreenath SN, Weis MC, & Stefan T (2012) Dynamic epitope expression from static cytometry data: principles and reproducibility. *PLoS ONE* 7(2):e30870.

19. King CL, *et al.* (2011) Fy(a)/Fy(b) antigen polymorphism in human erythrocyte Duffy antigen affects susceptibility to Plasmodium vivax malaria. *Proc Natl Acad Sci U S A* 108(50):20113-20118.

20. Tran TM, *et al.* (2005) Detection of a Plasmodium vivax erythrocyte binding protein by flow cytometry. *Cytometry A* 63(1):59-66.

21. Chaudhuri A, *et al.* (1994) Expression of the Duffy antigen in K562 cells. Evidence that it is the human erythrocyte chemokine receptor. *J Biol Chem* 269(11):7835-7838.
